# Catalytic Use of a Leader Peptide in the Biosynthesis of 3-Thiaglutamate

**DOI:** 10.1101/338681

**Authors:** Michael A. Funk, Chi Ting, Wilfred A. van der Donk

## Abstract

Small molecule natural products are key modulators of many types of intra- and interspecies communication. The availability of genome sequences allows the discovery of pathways to previously unknown natural products. We describe here a pathway in which a ribosomally synthesized small peptide serves as a catalytic scaffold on which a small-molecule anti-metabolite is biosynthesized in *Pseudomonas syringae*. First, a cysteine residue is transferred from Cys-tRNA to the C-terminus of the peptide, a reaction that replaces ribosomal protein synthesis. Then, a translocation of the cysteine thiol from the β-carbon to the α-carbon is catalyzed by an oxidase that removes the β-carbon as formate. The resulting thiol is carboxymethylated and proteolysis releases 3-thiaglutamate, in the process regenerating the peptide scaffold. This pathway features three previously unknown biochemical processes.

---

Bioactive small molecules play important roles in communication, symbiosis, and competition^1^. Historically most of these natural products have been discovered by activity based screens, but an alternative avenue for their discovery starts with the identification of their biosynthetic gene clusters. The bacterial genomes have revealed the tremendous diversity of natural products that remain to be discovered^2^. In this study we focused on a group of biosynthetic gene clusters for which the final products were not known and could not be predicted. Through a combination of *in vitro* experiments with purified enzymes and heterologous expression we show that these clusters encode an unusual biosynthetic pathway towards a glutamate analog that takes place on a scaffold peptide in the plant pathogen *Pseudomonas syringae* pv. maculicola ES4326.

Ribosomally synthesized and posttranslationally modified peptides (RiPPs) are a large and diverse class of natural products^3^. Lantibiotics and thiopeptides are two examples of RiPPs with various societal uses^4^. They are biosynthesized from a precursor peptide consisting of a leader peptide that serves as a recognition motif for the biosynthetic enzymes, and a core peptide that is converted to the final product. During the maturation of both classes of compounds, Ser and Thr residues are glutamylated by LanB enzymes in a glutamyl-tRNA dependent mechanism^5, 6^. Subsequently, the glutamate is eliminated to generate dehydroamino acids (Fig. 1a). A survey of the available genomes revealed a large number of LanB-like genes in which the domain responsible for glutamate elimination is missing, suggesting that these small LanBs might add an amino acid in a tRNA-dependent fashion without subsequent elimination.

**Figure 1.**
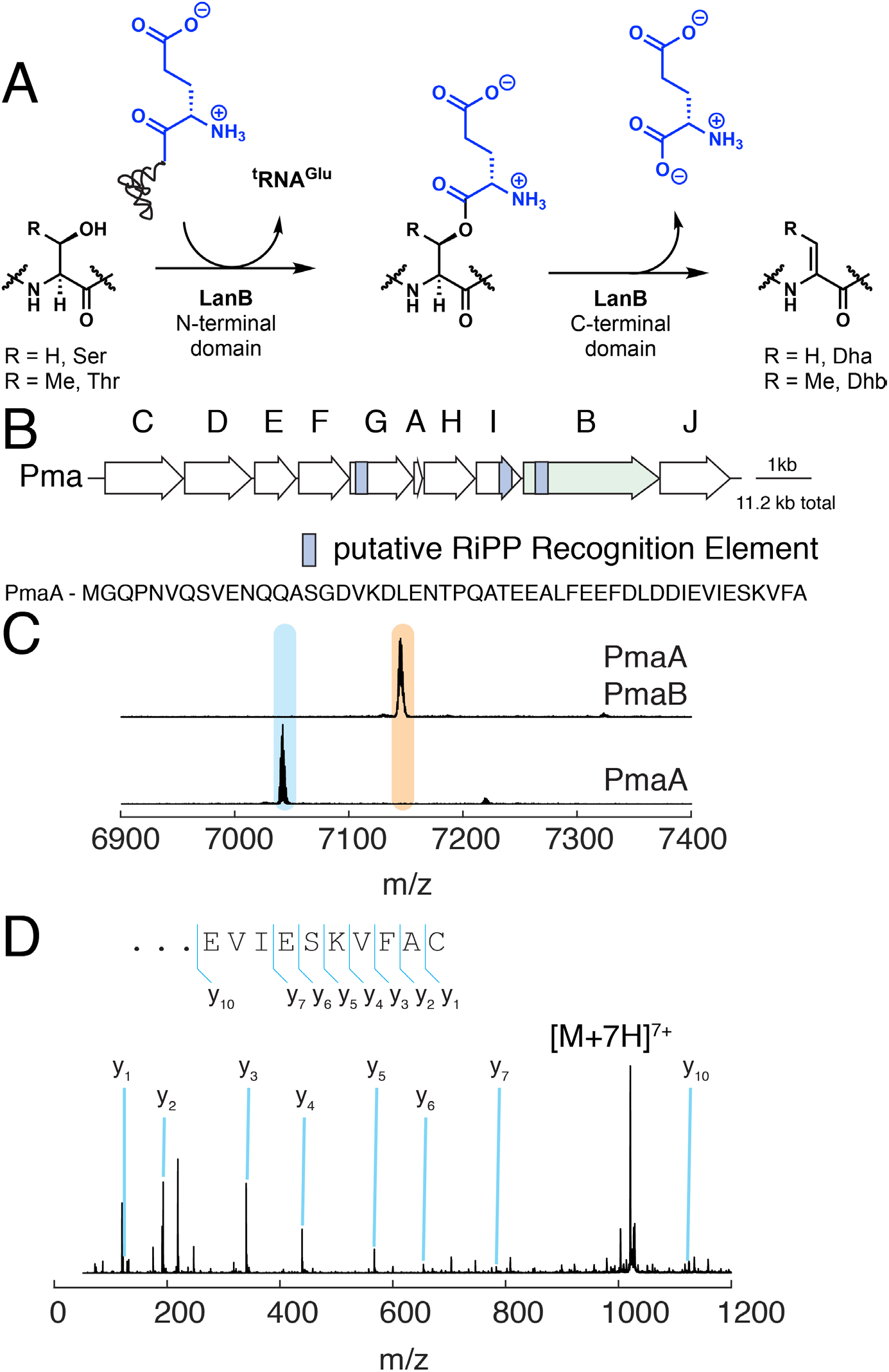
Function of a small LanB enzyme, PmaB, found in *P. syringae*. **a**, LanB enzymes glutamylate Ser/Thr residues and subsequently eliminate the glutamate to form dehydroamino acids. Small LanB proteins lack the elimination domain. Dha, dehydroalanine; Dhb, dehydrobutyrine. **b**, Biosynthetic gene cluster in *P. syringae* encoding a small LanB. **c**, Matrix-assisted laser desorption ionization with time-of-flight (MALDI-TOF) mass spectra of PmaA coexpressed with PmaB. **d**, Analysis of the PmaB product by tandem electrospray ionization (ESI) mass spectrometry.

In *P. syringae* such a small LanB (PmaB) is encoded near an open reading frame encoding a 42 amino acid peptide (PmaA; Fig. 1b). No Cys/Ser/Thr-rich core peptide is apparent in PmaA, but it does contain a region containing acidic and hydrophobic residues similar to other RiPP precursor peptides^7^. Co-expression of His_6_-PmaA and PmaB in *Escherichia coli* and subsequent affinity purification of the peptide demonstrated an increase in mass by 103 Da (Fig. 1c). This increase is not consistent with glutamylation, but could be the result of condensation with a cysteine residue. Glutamylation of the side chain of Ser/Thr results in an adduct that can be readily removed under basic conditions^8^, but such treatment of the peptide produced by PmaB did not lead to any hydrolysis (Extended Data Fig. 1). Subsequent high resolution mass spectrometry analysis of the peptide suggested that the adduct was unexpectedly attached to the C-terminal alanine (Fig. 1d), a linkage that could only be formed by a terminal amide. We expressed PmaA and PmaB individually as His_6_-tagged proteins and purified them. In vitro incubation with Cys, ATP, tRNA^Cys^ and Cys tRNA synthetase (CysRS) resulted in the same product (PmaA-Cys; Fig. 2a) as isolated from co-expression in *E. coli* confirming that PmaB adds a Cys to the C-terminus of PmaA in a tRNA dependent manner (Extended Data Fig. 2). This finding was unexpected as it constitutes a previously unknown type of posttranslational modification and seems counterintuitive since a more logical route to the product appears to be encoding the Cys on *pmaA*.

**Figure 2.**
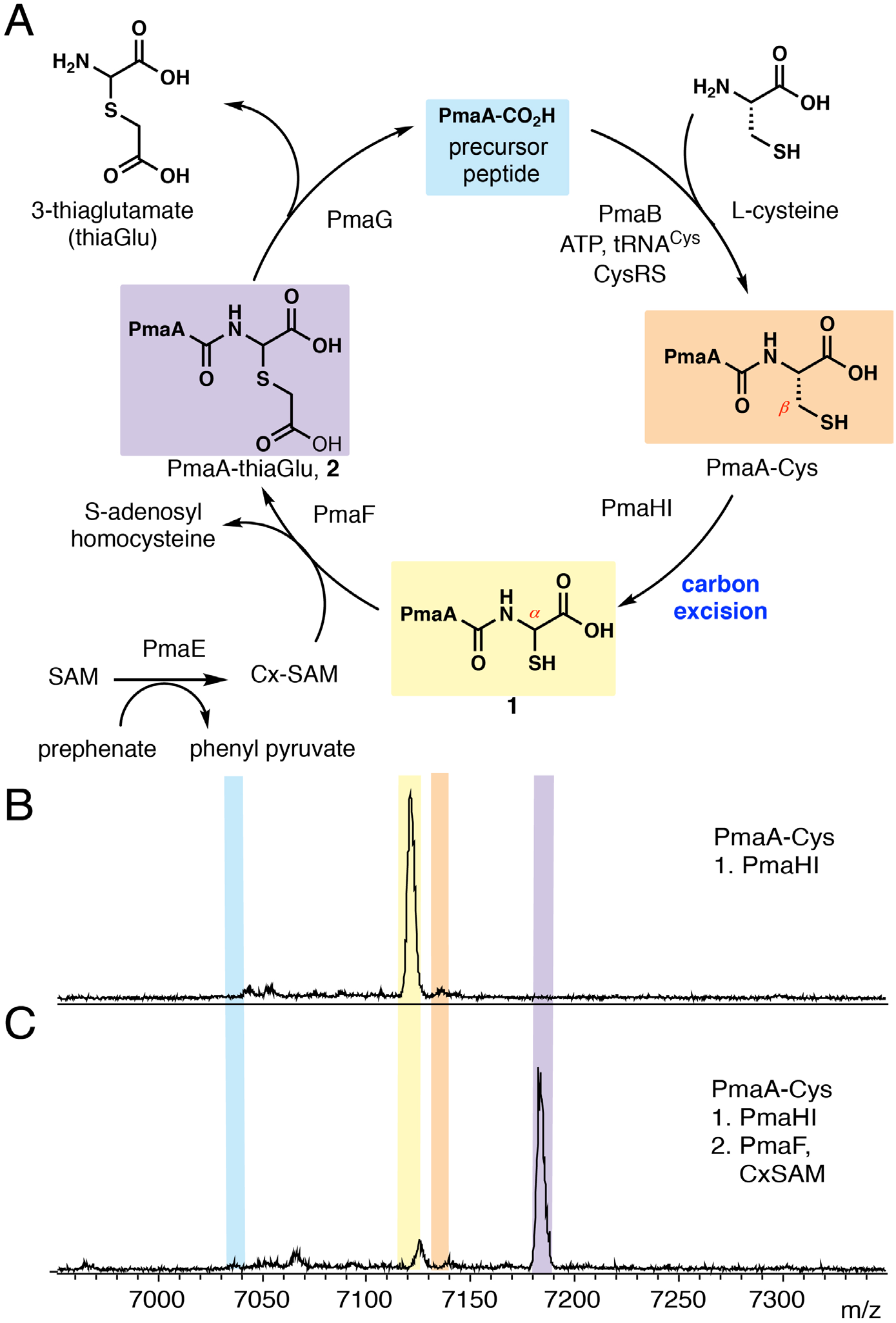
The cysteine added by PmaB is modified by other enzymes from the *pma* cluster. **a**, Inferred biosynthetic pathway towards 3–thiaglutamate. **b**, MALDI-TOF mass spectrum of PmaHI *in vitro* reaction with PmaA-Cys. **c**, MALDI-TOF mass spectrum of PmaF *in vitro* reaction with compound **1**. Color-coding of shaded peaks in panels B and C are shown in panel A.

Intrigued by the unexpected chemistry catalyzed by PmaB, we interrogated the other proteins encoded in the biosynthetic gene cluster. PmaH has low level homology to a dinuclear non-heme iron dependent protein for which a structure is known (PDB 3BWW) but for which no activity is reported. The C-terminal domain of PmaI has homology with known leader peptide binding domains in RiPP biosynthetic enzymes (Fig. 1b)^5, 9^ We co-expressed PmaA with PmaB, PmaH and PmaI in *E. coli* and isolated a product that was decreased in mass by 14 Da from PmaA-Cys (Extended Data Fig. 3). We treated the peptide with trypsin to generate a C-terminal tetrapeptide. Chemical assays with thiol-reactive and carboxylate-reactive electrophiles indicated that the product still contained a thiol group and a carboxylate (Extended Data Fig. 4) suggesting structure **1** as the product of PmaHI (Fig. 2a). We next repeated this experiment but using an *E. coli* strain that is auxotrophic for Cys and that was grown in minimal media supplemented with ^13^C-labeled Cys. Isolation of the peptide and analysis by MS showed that it is the cysteine β-carbon that is removed (Extended Data Fig. 5).

We noted peaks in the MS that were of intermediate mass between the native PmaA and the cysteine-modified peptide (Extended Data Fig. 3), suggesting that structure **1** is not stable. Analysis of the biosynthetic cluster revealed a pair of genes (*pmaEF*) encoding proteins similar to a recently characterized carboxy-*S*-adenosylmethionine (Cx-SAM) synthase and a SAM-dependent methyltransferase, respectively^10, 11^. We therefore subjected compound **1** to Cx-SAM and PmaF *in vitro* and isolated a new product with a mass increase of 58 Da (Fig. 2c). This increase is consistent with carboxymethylation of a thiol in the peptide by PmaF, which was confirmed by treating the PmaHI product with iodoacetic acid, which resulted in the same product. The peptide was treated with trypsin and the C-terminal tetrapeptide product characterized by ^1^H NMR spectroscopy and tandem MS, providing spectra consistent with structure **2** (Fig. 2a; Extended Data Fig. 6). Given the highly unusual architecture, we also chemically synthesized the peptide as two diastereomers (Extended Data Fig. 7) and demonstrated that the ^1^H NMR spectrum of one isomer was identical to the enzymatic product (Extended Data Fig. 6). Thus, collectively PmaBEFHI convert PmaA into a peptide containing a 3-thiaglutamate at its C-terminus (PmaA-thiaGlu, **2**; Fig. 2a).

Intrigued by the highly unusual transformation that appears to be catalyzed by PmaHI, we investigated this process with purified proteins. Neither protein could be expressed in soluble form individually, but co-expression of His_6_-PmaH with non-tagged PmaI and subsequent affinity chromatography resulted in co-purification of the proteins. Metal analysis indicated the presence of two Fe per PmaHI. *In vitro* PmaHI converted PmaA-Cys to **1** under aerobic conditions (Fig. 2b). However under low oxygen concentrations, product formation was negligible confirming the oxygen-dependency of the reaction (Extended Data Fig. 8). To investigate if PmaHI can functionalize internal cysteine residues, an insertion mutant, *pmaA-CysAla*, was prepared. The expressed peptide, PmaA-CysAla, was not modified by PmaHI, signifying that a C-terminal cysteine is required for the reaction to occur (Extended Data Fig. 9). In order to identify the fate of the lost carbon atom, ^13^C-labeled PmaA-Cys was expressed in the *E. coli* cysteine auxotroph using 3-^13^C-cysteine. When ^13^C-labeled PmaA-Cys was reacted with PmaHI, formate was observed as a product by ^13^C NMR (Fig. 3). Thus, PmaHI catalyzes a net four-electron oxidation of PmaA-Cys, modifying the redox states of both the α and β carbons of the C-terminal cysteine installed by the small LanB enzyme.

**Figure 3.**
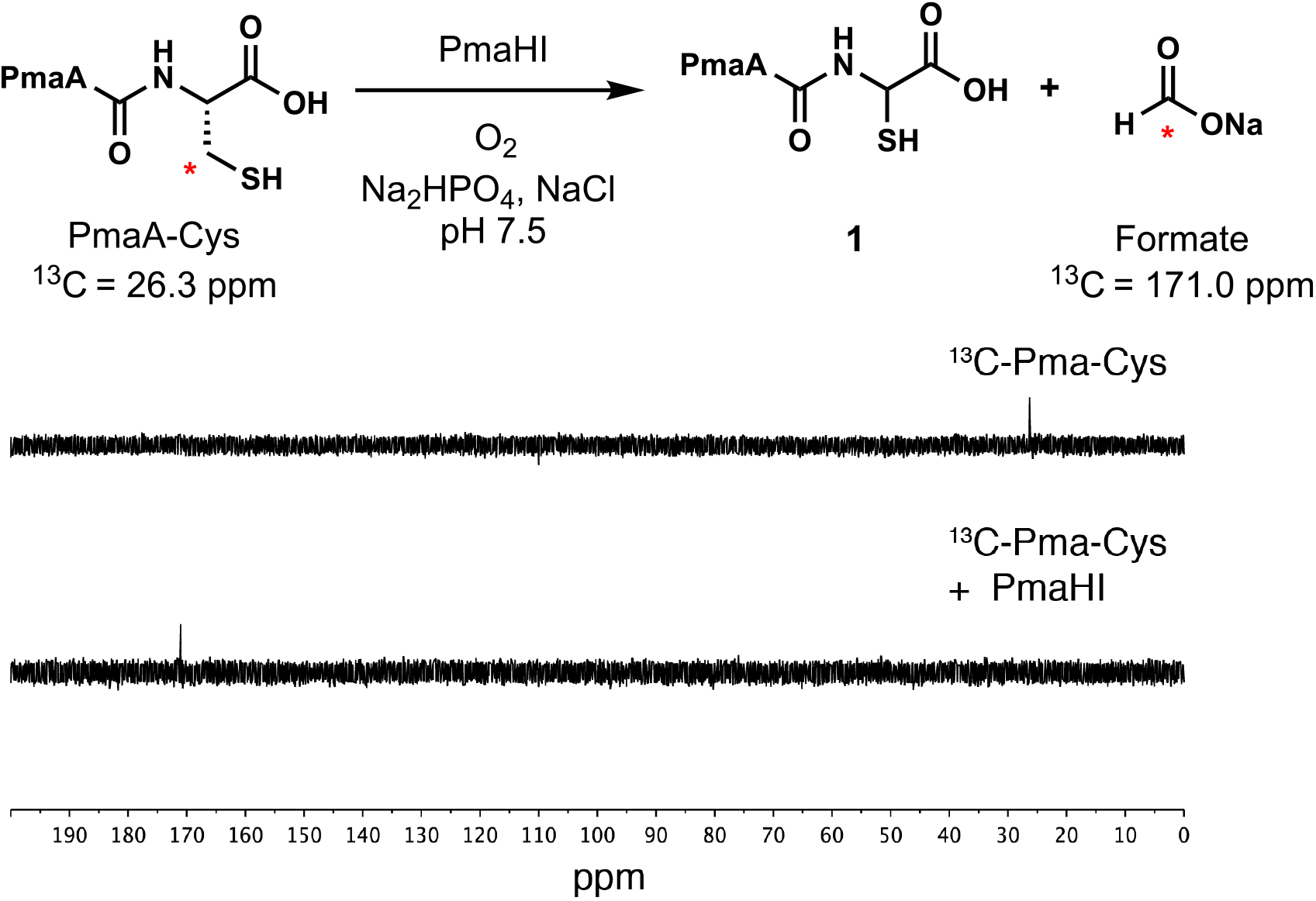
*In vitro* PmaHI reacts with ^13^C-labeled PmaA-Cys to produce ^13^C-formate. The signal at 26.3 ppm in the ^13^C NMR spectrum corresponds to the *β*-carbon of the C-terminal cysteine in PmaA-Cys. After reaction with PmaHI, a new signal at 171.0 ppm is observed that corresponds to ^13^C-formate.

Having identified the products of the PmaHI reaction, we propose a mechanism for the formation of **1** and formate from PmaA-Cys. Because PmaA-CysAla was not a substrate, the C-terminal carboxylate of PmaA-Cys is invoked for substrate binding to the diiron active site. Similar to the non-heme diiron enzyme *myo*-inositol oxygenase (MIOX)^12, 13^, the reaction is likely initiated via hydrogen atom abstraction by a Fe^III^-superoxo species from substrate, in this case the *β*-carbon of cysteine (**3** → **4**, Fig. 4). Hydroxylation by the Fe^III^-hydroperoxo intermediate would produce a thioacetal bound to a ferryl (Fe^IV^=O) complex (**4** → **5**, Fig. 4). Carbon-carbon bond fragmentation would be triggered by hydrogen atom abstraction from the hydroxyl group by the ferryl intermediate to generate an oxygen radical which undergoes *β*-scission (**5** → **6** → **7**, Fig. 4). This process bares resemblance to a step in the reaction catalyzed by 2-hydroxyethylphosphonate dioxygenase (HEPD)^14^. Sulfur rebound of the thioformate ligand forges the carbon-sulfur bond in **1** and regenerates the Fe^II^ complex, a step with similarity to thioether formation by isopenicillin N synthase^15^. Finally, hydrolysis of the thioformyl group by the hydroxide ligand on iron produces both formate and **1** after ligand exchange, effectively closing the catalytic cycle (Fig. 4).

**Figure 4.**
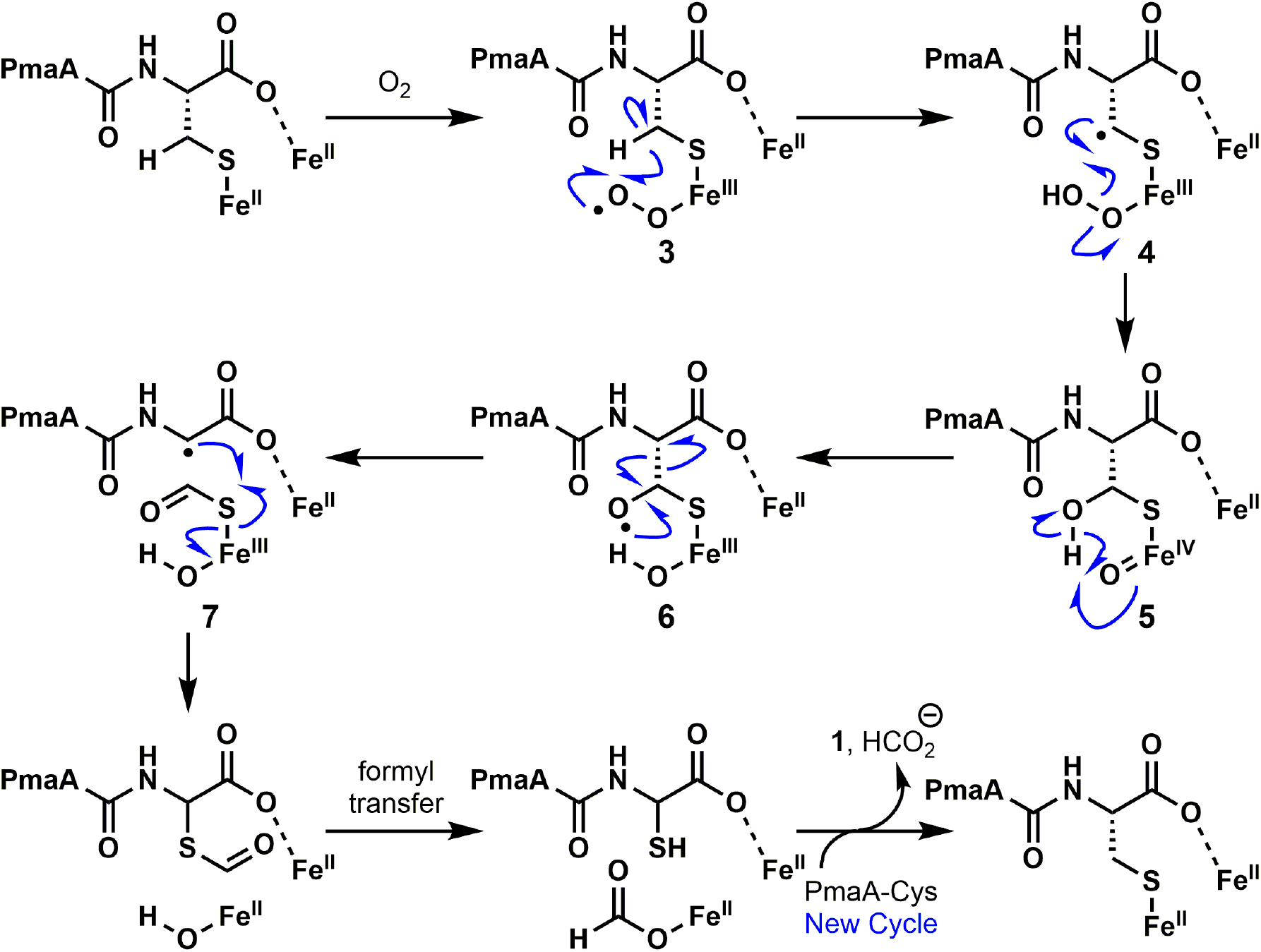
Proposed mechanism for the carbon excision reaction catalyzed by PmaHI. The two iron atoms might be connected by bridging ligands. Conversion of **4** to **5** could also involve electron transfer from the radical to the Fe(III), thus generating a thioaldehyde and a Fe(II)-hydroperoxo. The latter could expel hydroxide and form a ferryl, and the hydroxide could hydrate the thioaldehyde, arriving again at intermediate **5**. A related transformation is invoked in HEPD^14^.

The last four genes in the biosynthetic cluster encode a putative membrane bound protease (PmaG), a putative pyridoxal-phosphate dependent enzyme (PmaC), and two putative transporters (PmaD and PmaJ). Like PmaB and PmaI, PmaG also contains a RiPP leader peptide recognition motif suggesting it will act on the PmaA-derived peptide. (Fig. 1b). When PmaA-thiaGlu was exposed to the membrane fraction of cell lysate of *E. coli* expressing GFP-PmaG, the peptide was cleanly converted into the starting PmaA peptide (Extended Data Fig. 10). PmaA-Glu was also a substrate but not PmaA-GluAlaa illustrating that the protease cannot distinguish Glu and 3-thiaGlu, but does not tolerate extension of the peptide. Thus, PmaA appears to be a scaffold on which thiaGlu is assembled and final proteolytic release of thiaGlu regenerates PmaA for another round of biosynthesis (Fig. 2a). Were cysteine merely encoded in *pmaA*, then each ribosomally produced peptide could make only a single 3-thiaglutamate. Instead, the catalytic use of PmaA as a leader peptide uncovered here is more efficient than the stoichiometric use of leader peptide in other RiPP pathways^4^. Perhaps this unusual pathway evolved because of the significant relative burden of leader peptide production for a single amino acid product. Bioinformatic prediction of PmaA transcriptional regulation^16^ suggests precursor production does not appear to be driven by a separate promoter (Extended Data Fig. 11). This contrasts with most other RiPP pathways in which expression of the substrate peptide is controlled by its own promoter followed by a read-through transcriptional terminator to allow the precursor peptide to be present in excess over the biosynthetic machinery^17, 18^. It is the Cys-tRNA-dependent enzyme PmaB that allows the catalytic use of PmaA. All other enzymes could in principle, and do in our co-expression studies, act on a genetically encoded cysteine at the C-terminus of PmaA.

This study shows that the gene cluster in *P. syringae* governs the biosynthesis of a RiPP consisting of just a single amino acid. Unlike other RiPPs, the final product in this pathway is not encoded in the genome since the pathway does not contain the gene-encoded core peptide^4^. Amino acyl-tRNA dependent amide bond formation to the C-terminus of a peptide is complementary to peptide bond formation by the ribosome, which extends peptides at their N-terminus. Unlike the ribosomal process, the substrate peptide of PmaB is not tRNA-bound. Small LanB-encoding genes are abundant in bacterial genomes, with some clusters encoding multiple small LanB proteins (Extended Data Fig. 12). Thus, this mechanism of peptide bond formation appears to be widespread and is likely driven by the advantages offered by catalytic use of leader peptides. The chemistry described herein (Fig. 2a) expands the range of post-translational modifications in RiPP biosynthesis^19^ to include a remarkable excision of a methylene group from cysteine. Similar enzyme systems are present in the genomes (Extended Data Fig. 13) including in the biosynthetic gene cluster encoding the methanobactin precursor^20–22^. The observed biosynthesis of a metabolite on a small peptide scaffold is unusual, with the closest similarity found in the biosynthesis of amino acids linked by isopeptide bonds to a glutamate residue on amino-carrier proteins in some bacteria^23, 24^.

## Acknowledgements

This work was supported by the National Institutes of Health (R37 GM058822 to W.A.V.; F32 GM120868 to M.A.F.). W.A.V. is a Howard Hughes Medical Institute Investigator. *P. syringae* pv. maculicola ES4326 was generously provided by Prof. Darrell Desveaux (University of Toronto).

## Author contributions

M.A.F. and C.P.T. performed biochemical assays. M.A.F. performed bioinformatics analysis. M.A.F., C.P.T., and W.A.V. designed experiments, analyzed data, and wrote the manuscript. M.A.F. and C.P.T. contributed equally to this study.

## Author Information

The authors declare no competing financial interest. Correspondence and requests for materials should be addressed to W.A.V.

## Methods

### Figure preparation

Gene cluster diagrams (Fig. 1b, Extended Data Figure 12 and 13) were made with EasyFig^25^. Mass spectra were plotted in Matlab R2015b. All figures were constructed with Adobe Illustrator. NMR spectra were analyzed in Mnova (Mestrelab Research).

### Materials

Materials were obtained from Sigma-Aldrich unless otherwise noted. All water was deionized and purified using a Milli-Q IQ 7000 purifier. MALDI-TOF MS was performed on a Bruker Daltonics UltrafleXtreme MALDI TOF/TOF instrument. ESI-MS analyses were performed using a SYNAPT ESI quadrupole TOF Mass Spectrometry System (Waters) equipped with an ACQUITY Ultra Performance Liquid Chromatography (UPLC) system (Waters).

### Cloning and cell culture

DNA was prepared using MiniPrep kits (Qiagen) using the manufacturer’s instructions from *E. coli* DH10b cells (New England Biolabs) made chemically competent by the KCM method^26^. Genomic DNA from *P. syringae* pv. maculicola ES4326 (Pma) was prepared using an UltraClean Microbial DNA isolation kit (Mo Bio Laboratories) according to the manufacturer’s instructions; Pma cells were grown in liquid KB medium with vigorous aeration. Plasmids were constructed with type-II restriction enzymes using the New England Biolabs Golden Gate Assembly Tool (http://goldengate.neb.com/editor) or manually designed and prepared by Gibson assembly. Restriction enzymes and PCR polymerases were obtained from New England Biolabs. Genes for each of the *pma* gene cluster products and the *P. syringae* cysteine tRNA synthetase (CysRS) were identified using the Joint Genome Initiative Integrated Microbial Genomes and Microbiomes webtool (http://img.jgi.doe.gov). Primers for cloning and site-directed mutagenesis were obtained from Integrated DNA Technologies. Mutagenesis was accomplished using QuikChange II Site-Directed Mutagenesis Kit according to manufacturer’s instructions. Primers are listed in Extended Data Table 1.

### Peptide expression and purification

For peptide expression, *E. coli* BL21(DE3) cells (New England Biolabs) expressing N-terminal His_6_-PmaA from pRSF-Duet were grown with 50 μg/mL kanamycin in homemade autoinduction (AI) media (for 1 L total: 10 g bacto tryptone, 5 g yeast extract, 5 g NaCl, 3 g KH_2_PO_4_, 6 g Na_2_HPO_4_) with 1X AI sugar solution containing 0.5% (vol/vol) glycerol, 0.05% (w/vol) glucose, and 0.2% (w/vol) lactose (20 mL of 50X stock). Cultures were shaken at 37 °C for 6-7 h following inoculation with 1 mL saturated culture. Cells were harvested and lysed by sonication. Peptides were purified by immobilized metal affinity chromatography (IMAC). The lysate was clarified by centrifugation at 29,000 rcf and applied to a 5 mL NiNTA-agarose column (GE Healthcare) using a peristaltic pump. The immobilized peptide was washed with 5 column volumes (CV) of 90% lysis buffer (50 mM HEPES pH 7.5, 100 mM NaCl), 10% elution buffer (50 mM HEPES pH 7.5, 100 mM NaCl, 500 mM imidazole; final imidazole concentration in wash: 50 mM) and eluted with 100% elution buffer. The elution fraction was concentrated using a 3 kDa MWCO Amicon spin filter and washed with 10-20 CV of deionized water to remove imidazole. Crude peptide was desalted with a VYDAC^®^ Bioselect C4 cartridge. Peptide elution fractions were lyophilized.

His_6_-PmaA-Cys and the modified peptide **1** obtained by co-expression of PmaA with PmaH/I/B were obtained using the same procedure described above. For analysis of modified PmaA peptides, the same procedure was used but the peptide was reacted with 20 mM N-ethylmaleimide (NEM) or 20 mM 2-iodoacetic acid (IAA) prior to concentration by Amicon. To reduce the size of the peptide, the lyophilized product was resolubilized in denaturing buffer (1 mL 6 M guanidinium chloride, 20 mM NaH_2_PO_4_, 500 mM NaCL, 0.5 M imidazole, pH 7.5), diluted 10-fold and digested by trypsin (0.5 mg/mL) to release the VFAC-NEM, VFAX-NEM (X is the thioaminal in structure **1**), or VFA-thiaGlu fragments.

### Protein expression and purification for PmaB, PmaF and CysRS

All heterologously-expressed proteins were obtained from *E. coli* BL21(DE3) cells (New England Biolabs) made chemically competent through the KCM method^26^. The general expression protocol for the *pma* cluster enzymes PmaB, and PmaF is as follows: A 1 mL inoculum was added to 1 L of AI medium containing 50 μg/mL kanamycin and 1x AI sugars. For N-terminal His_6_-PmaB or His_6_-CysRS, cells were grown LB medium and expression was induced with 0.5 μM isopropyl-beta-D-1-thiogalactopyranoside. Culture were grown with vigorous shaking at 37 °C for 3-4 h and then shifted to 21 °C for overnight (~10 h) expression. Cells were harvested by centrifugation, collected in 50 mL tubes and frozen in liquid nitrogen. For purification of His_6_-PmaB and His_6_-CysRS, cells were thawed, resuspended in lysis buffer (50 mM HEPES pH 7.5, 100 mM NaCl; 30 mL per 10 g wet cell paste) and lysed by treatment with lysozyme (100 μg/mL) and sonication (3 min active time; 1 s pulse, 2 s rest at 60 % max amplitude using a 1 cm tip). Proteins were purified by immobilized metal affinity chromatography (IMAC). Phenylmethylsulfonyl fluoride was added to prevent degradation by serine proteases and 1 mM TCEP was included to maintain reduced thiols. The eluate was concentrated to 2.5 mL in a 30 kDa MWCO centrifuge filter and desalted on a PD-10 size-exclusion column (GE Healthcare Life Sciences). Protein was separated into aliquots and stored at −78 °C. Protein purity was judged by SDS-PAGE (Extended Data Figure 14).

### Protein expression and purification for PmaHI

PmaHI was obtained from electrocompetent *E. coli* BL21(DE3) cells (New England Biolabs) transformed with pET15b-*pmaHI*. A 25 mL inoculum was added to 1 L of LB medium containing 100 μg/mL ampicillin. Cells were grown with vigorous shaking at 37 °C for 3-4 h until the optical density (OD) at 600 nm reached 0.6 at which point the culture was cooled to 0 °C and induced with 1 mM isopropyl-beta-D-1-thiogalactopyranoside. Upon induction, the cell culture was incubated at 18 °C for 14 h with vigorous shaking. Cells were harvested by centrifugation, collected in 50 mL tubes and suspended in lysis buffer (20 mM Tris, 300 mM NaCl, 10% glycerol, pH 7.6,) supplemented with 1 mg/mL lysozyme, 600 U DNase, and 1 mM TCEP. The cells were lysed by passage through a French pressure cell twice and cell debris was removed by centrifugation (30,000 rcf) for 50 min at 10 °C. The supernatant was loaded onto a column containing 5 mL of Ni-NTA resin previously equilibrated with lysis buffer. After equilibration of the resin with the lysate by orbiting for 30 min, the flow-through was discarded. The resin was washed with 2 × 40 mL of wash buffer (25 mM imidazole, 20 mM Tris, 300 mM NaCl, 10% glycerol, pH 7.6,) followed by elution with elution buffer (250 mM imidazole, 20 mM Tris, 300 mM NaCl, 10% glycerol, pH 7.6,). The eluate was concentrated to 2.5 mL in a 30 kDa MWCO centrifuge filter and desalted on a PD-10 size-exclusion column (GE Healthcare Life Sciences). Protein was separated into aliquots and stored at −78 °C. The protein was judged pure by SDS-PAGE (Extended Data Figure 14).

### HPLC peptide purification and LC-MS

Following IMAC and C4 desalting, PmaA was purified using an Agilent 1200 HPLC. HPLC buffers were 0.1% TFA in water (HPLC buffer A) and 0.1% TFA in acetonitrile (HPLC buffer B). The desalted material was lyophilized, resuspended in buffer A, and applied to a Phenomenenx Luna C18 column equilibrated with 95% HPLC buffer A, 5% HPLC buffer B. A linear gradient to 100% HPLC buffer B was run over 20 min at 0.1 mL/min. PmaA eluted near the end of the gradient. PmaA-Cys behaved similarly and eluted slightly later. Full-length and AspN or trypsin-digested peptides were analyzed by LC-MS/MS on a Waters Synapt Q-TOF equipped with a Waters UPLC and Phenomenex C18 Luna or C4 Jupiter columns. Buffers for LC-MS were 0.1% formic acid in water (LC-MS buffer A) or 0.1% formic acid in LC-MS grade acetonitrile (LC-MS buffer B).

### Analysis of metal concentration

Iron quantification of PmaHI was determined using Ferene S as a spectrophotometric dye as reported by Hennessy and co-workers^27^. A standard curve was generated using an iron standard in 2% HNO3 solution (Claritas PPT).

### Chemical synthesis of CxSAM

CxSAM was synthesized and purified as previously described^10^. In a 1.5 mL tube, S-adenosylhomocysteine (3 mg) and iodoacetic acid (0.1 g) were dissolved in 150 mM ammonium bicarbonate (0.5 mL) and incubated at 37 °C for 12 h with shaking. The resulting mixture was analyzed by LC-MS and purified by analytical C18 HPLC. CxSAM is poorly retained on C18 and eluted from the column at 5 min in 0% HPLC buffer B (~3.5 min dead time). HPLC fractions were analyzed by LC-MS and had characteristic fragments for S-carboxymethyl thioadenosine.

### Attempted hydrolysis of PmaA-Cys

PmaA-Cys was obtained by co-expression of His_6_-PmaA and PmaB in *E. coli* and purified by the methods described above. PmaA-Cys (100 μM) in HEPES buffer (50 mM HEPES, 100 mM NaCl, pH 7.5) was basified with 1 M NaOH until pH 12 and incubated at room temperature for 1 h. The reaction mixture was then directly desalted and concentrated by ZipTip, eluted in 80% acetonitrile, 20% water, 0.1% TFA, and analyzed by MALDI-TOF MS (Extended Data Figure 1).

### In vitro assay of PmaHI

Carbon excision of PmaA-Cys by PmaHI was monitored by MALDI-TOF MS using parameters similar to those described above. In a 2.5 mL centrifuge tube, PmaA-Cys (100 μM) was added to PmaHI (10 μM) in phosphate buffer (0.2 mL, 50 mM Na_2_HPO_4_, 300 mM NaCl, 10% glycerol, pH 7.6). The reaction vessel was left open to air at 24 °C for 16 h. At this time, the reaction was directly desalted and concentrated by ZipTip, eluted in 80% acetonitrile, 20% water, 0.1% TFA, and analyzed by MALDI-TOF MS (Fig. 2b). PmaA-CysAla was subjected to identical reaction conditions as mentioned above. For low oxygen reactions, PmaHI was subjected to buffer exchange with degassed phosphate buffer 10 times using a 10 kDa MWCO Amicon 0.5 mL spin filter in an anaerobic chamber maintained at < 1.0 ppm oxygen. PmaA-Cys was also subjected to buffer exchange with degassed phosphate buffer using a 3 kDa MWCO Amicon 0.5 mL spin filter in the anaerobic chamber. PmaA-Cys (100 μM) was added to PmaHI (10 μM) in phosphate buffer (0.2 mL, 50 mM Na_2_HPO_4_, 300 mM NaCl, 10% glycerol, pH 7.6) and incubated in the anaerobic chamber for 1 h. At this point, the enzyme was first deactivated with 1 M formic acid before exposing the reaction mixture to air. The reaction mixture was then analyzed as described above.

### In vitro assay of PmaF

Carboxymethylation of compound **1** by PmaF was monitored by MALDI-TOF and LC-MS using parameters similar to those described above. In a 2.5 mL centrifuge tube, peptide **1** (40 μM) and Cx-SAM (120 μM) was added to PmaF (10 μM) in HEPES buffer (0.5 mL, 50 mM HEPES, 100 mM NaCl, pH 7.6). After 13 h, the reaction was directly desalted and concentrated by ZipTip, eluted in 80% acetonitrile, 20% water, 0.1% TFA, and analyzed by MALDI-MS (Fig. 2c).

### Chemical analysis of modified PmaA peptide

PmaBHI-modified PmaA was first treated with trypsin as described under *Peptide expression and purification*. Peptide fragments were esterified using methanolic HCl^28^. First 160 μL of acetyl chloride was added dropwise to 1 mL of anhydrous methanol while stirring in an ice bath. Then 100 μL of this mixture was added to dry peptide and incubated at room temperature for 4 h. The esterification mixture was diluted with 400 μL of water, frozen in liquid nitrogen, and lyophilized. The modified peptides were analyzed by LC-MS as described above.

### Proteolytic activity of GFP-PmaG membrane fraction

For protein expression of GFP-PmaG, *E. coli* BL21(DE3) cells (New England Biolabs) transformed with pet28b-GFP-*pmaG* were grown with 50 μg/mL kanamycin in LB media (for 1 L total: 10 g bacto tryptone, 5 g yeast extract, 10 g NaCl). A 25 mL starting culture was inoculated with a single colony and shaken at 37 °C for 14 h followed by 40x dilution into 1 L of LB containing 50 μg/mL kanamycin. The culture was kept at 37 °C until optical density at 600 nm reached 0.6-0.8. Then, the flask was placed in an ice/water bath for 30 min before the addition of isopropylthio-β-D-galactoside (IPTG) to a final concentration of 1 mM. The culture was then incubated for an additional 14 h at 18 °C. At this time, the cells were collected by centrifugation at 11,270 rcf for 20 min. Cells were harvested, resuspended in 25 mL of Tris lysis buffer (20 mM Tris, 300 mM NaCl, 10 mM imidazole, 10% glycerol, pH 7.6), and 10 mg lysozyme, TCEP (1 mM) and 600 U DNAse were added. Cells were lysed by sonication, and the cell lysate (0.4 mL) was added to PmaA-thiaGlu (**2**, prepared separately) in 0.1 mL HEPES buffer (50 mM HEPES, 100 mM NaCl, pH 7.6) at rt. After 16 h, the reaction was directly desalted and concentrated by ZipTip, eluted in 80% acetonitrile, 20% water, 0.1% TFA, and analyzed by MALDI-MS (Extended Data Figure 10). PmaA-Glu and PmaA-GluAla were subjected to identical reaction conditions as mentioned above.

### Isotope Labeling Experiments

^13^C-cysteine labeled His_6_-PmaA-Cys was expressed using *E. coli* strain JW3582-2. *E. coli* JW3582-2, an auxotrophic strain for cysteine, was purchased from the Coli Genetic Stock Center at Yale University, https://cgsc2.biology.yale.edu/Strain.php?ID=108920. A lysogenization step was performed to JW3582-2 so that the host strain could be used to express target genes cloned in T7 expression vectors. Lysogenization was performed using the λDE3 Lysogenization Kit (Novagen) as per the manufacturer’s instructions. The lysogenized JW3582-2 was transformed with a pACYCDuet-1 plasmid encoding His_6_-PmaA-Cys, PmaH and PmaI. Expression of ^13^C labeled, modified His_6_-PmaA-Cys was performed in modified M9 minimal media. A starter culture of *E. coli* BL21(DE3) cells containing a pACYCDuet-1 plasmid encoding His_6_-PmaA-Cys, PmaH and PmaI were grown overnight at 37 °C in lysogeny broth (LB) containing 25 μg/mL kanamycin and 20 μg/mL chloramphenicol. After harvesting the cells, the supernatant was discarded and the cells were washed with 5 mL of wash buffer (22 mM KH_2_PO_4_, 42 mM Na_2_HPO_4_, 8.5 mM NaCl, pH 7.4). After washing, the cells were resuspended in wash buffer and used to inoculate (1:200) modified M9 minimal media, with the following composition per 100 mL:10 mL of a 10x minimal media (220 mM KH_2_PO_4_, 420 mM Na_2_HPO_4_, 85 mM NaCl, pH 7.4), 0.3 mL of 40% aqueous (NH_4_)_2_SO_4_, 2 mL of 20% aqueous glucose, 0.1 mg of FeSO_4_, 10 μg of thiamine, 200 μL of 1 M MgSO_4_, 10 μL of 1 M CaCl_2_, and 75 μL of a trace element solution (5 mM CaCl_2_, 1.25 mM ZnCl_2_, 260 μM CuCl_2_•H_2_O, 252 μM CoCl_2_•6H_2_O, 250 μM Na_2_MoO_4_•2H_2_O, pH 7.4). L-cysteine or U-^13^C-cysteine or 3-^13^C-cysteine (1 mM) was added as the sole cysteine source. The media also contained 10 μg/mL chloramphenicol and 25 μg/mL kanamycin. The cells were grown at 37 °C and induced at OD_600_ = 0.6-0.8 by the addition of IPTG to a final concentration of **1** mM and grown for an additional 3 h at 37 °C before harvesting. Peptide 1 was purified as described above. After modification with iodoacetic acid (20 mM) and trypsin digestion, the VFAX-CH_2_CO_2_H tetrapeptide was analyzed by LC/MS (Extended Data Figure 5). For preparation of ^13^C-enriched PmaA-Cys for *in vitro* PmaHI reaction and ^13^C NMR spectroscopy (Fig. 3), the above procedure was modified to use pACYC-Duet1 plasmid encoding only His_6_-PmaA-Cys, and in the purification phosphate buffer (50 mM Na_2_HPO_4_, 300 mM NaCl, pH 7.5) was used instead of HEPES.

### Bioinformatics for operon prediction

The operon containing the *pma* gene cluster was predicted by the Softberry FGENESB program (Softberry, Inc., Mount Kisco, NY) (http://www.softberry.com/)^29^.

### General procedures for chemical synthesis

Unless stated otherwise, all reactions were performed in flame-dried glassware under an atmosphere of dry nitrogen or argon. Dry dichloromethane, and dimethylformamide (DMF) were obtained by passing these degassed solvents through activated alumina columns. All other reagents were used as received from commercial sources, unless stated otherwise. Reactions were monitored by thin layer chromatography (TLC) on Silicycle Siliaplate glass-backed TLC plates (250 μm thickness, 60 Å porosity, F-254 indicator) and visualized by UV irradiation or development with an anisaldehyde or phosphomolybdic/cerium sulfate stain. Volatile solvents were removed under reduced pressure with a rotary evaporator. All flash column chromatography was performed using Silicycle SiliaFlash F60, 230-400 mesh silica gel (40-63 μm).

^1^H NMR and ^13^C NMR spectra were recorded on an Agilent 600 MHz spectrometer for ^1^H (150 MHz for ^13^C) in D2O. Chemical shifts are reported relative to the residual solvent signal (^1^H NMR: δ = 4.79). NMR data are reported as follows: chemical shift (multiplicity, coupling constants where applicable, number of hydrogens). Splitting is reported with the following symbols: s = singlet, bs = broad singlet, d = doublet, t = triplet, app t = apparent triplet, dd = doublet of doublets, ddd = doublet of doublet of doublets, dt = doublet of triplets, hept = heptet, m = multiplet. Infrared spectra (IR) were recorded as a thin film on a Perkin-Elmer FT-IR system and peaks were reported in cm^-1^.

#### Hemiaminal S1

**S1** was prepared on multi-gram scale according to the procedure reported by Rivier and co-workers^30^. An oven-dried 250 mL round-bottom flask was charged with glyoxylic acid monohydrate (3.6 g, 39 mmol, 1.1 equiv) and *tert*-butyl carbamate (4.1 g, 35 mmol, 1.0 equiv). The reaction vessel was evacuated and backfilled with nitrogen and this process repeated for a total of three times. Diethyl ether (25 mL) was added and the reaction mixture was stirred for 14 h at rt. The reaction mixture was filtered and concentrated *in vacuo*. The crude residue was dissolved in EtOAc and triturated with hexanes to afford **S1** (4.9 g, 73% yield) as a yellow-brown solid. Spectral data were in agreement with that previously reported by Rivier and co-workers^30^.

#### Ester S2

A 250 mL round-bottom flask equipped with a Dean-Stark trap and reflux condenser was charged with thioglycolic acid (3.0 g, 33 mmol, 1.0 equiv), benzyl alcohol (5.3 g, 49 mmol, 1.5 equiv) and *p*-toluenesulfonic acid monohydrate (0.63 g, 3.3 mmol, 0.1 equiv). The reaction vessel was evacuated and backfilled with nitrogen and this process repeated for a total of three times. Toluene (100 mL) was added and the reaction mixture was heated to reflux and maintained at this temperature for 14 h. The reaction mixture was cooled to rt, and then concentrated *in vacuo*. The crude residue was purified by column chromatography (5% → 7.5% EtOAc in hexanes) to afford ester **S2** (2.6 g, 43% yield) as a colorless oil. Spectra data were in agreement with that previously reported^31^.

#### Thioether S3

A 100 mL round-bottom flask equipped with a Dean-Stark trap and reflux condenser was charged with hemiaminal **S1** (1.0 g, 5.0 mmol, 1.0 equiv), ester **S2** (1.0 g, 5.5 mmol, 1.1 equiv) and *p*-toluenesulfonic acid monohydrate (60 mg, 0.31 mmol, 0.06 equiv). The reaction vessel was evacuated and backfilled with nitrogen and this process repeated for a total of three times. Toluene (100 mL) was added and the reaction mixture was heated to reflux and maintained at this temperature for 6 h. The reaction mixture was cooled to rt, and then concentrated *in vacuo*. The crude residue was purified by column chromatography (10% → 20% MeOH in CH_2_Cl_2_) to afford thioether **S3** (400 mg, 23% yield) as a brown oil: ^1^H NMR (600 MHz, D2O) δ 7.52 – 7.42 (m, 5H), 5.26 (d, J = 12.3 Hz, 1H), 5.23 (d, J = 12.3 Hz, 1H), 5.18 (bs, 1H), 3.52 (bs, 2H), 1.43 (s, 9H)); ^13^C NMR (150 MHz, D_2_O) δ 173.3, 172.5, 156.1, 135.2, 128.8, 128.6, 128.2, 81.6, 67.8, 58.8, 31.9, 27.5; IR (thin film) 3415, 3055, 2984, 1717, 1496, 1455 cm^-1^; HRMS (ESI) calcd for[C_16_H_22_NO_6_S]^+^ (M+H)^+^: m/z 356.1168 found 356.1156.

#### VFA-thiaGlu S4

**S3** was loaded on Merrifeld resin using the Gisin Method^32^. In a 20 mL scintillation vial, **S3** (350 mg, 1.0 mmol, 1 equiv) was dissolved in methanol (5 mL) and water (0.5 mL). The resulting solution was basified to pH 7.0 with 20% aqueous solution Cs_2_CO_3_ and then concentrated *in vacuo*. The residue solid was added DMF (2.5 mL) and then concentrated *in vacuo*, and this process was repeated a total of two times. In a separate 100 mL round-bottom flask, Merrifield resin (1.0 g, 1-1.5 mmol/g Cl^-^ loading, 2% cross linked) was added, followed by DMF (8 mL) and the mixture was gently stirred for 10 min. The cesium salt of **S3** was added to the flask containing the resin and the mixture was heated to 50 °C and maintained at that temperature for 18 h. The resin was filtered with a fritted glass funnel by vacuum filtration and washed successively with DMF, DMF/H_2_O (1:1), MeOH/H_2_O (1:1) and then MeOH. After the resin was dry, 50% TFA in CH_2_Cl_2_ (1 mL) was added to the resin on the fritted glass filter. After 3 min, the 50% TFA in CH_2_Cl_2_ was removed by vacuum filtration, and the resin was washed with CH_2_Cl_2_ (3 × 2 mL) followed by diisopropylethylamine (3 × 2 mL, 5% v/v in CH_2_Cl_2_). To the resin was then added Boc-L-Ala-OH (380 mg, 2.0 mmol, 2 equiv) in DMF (5 mL) followed by the addition of HOBt (270 mg, 2.0 mmol, 2 equiv), HBTU (760 mg, 2.0 mmol, 2 equiv), and diisopropylethylamine (0.7 mL, 4.0 mmol, 4.0 equiv) in DMF (2 mL). The reaction was mixed with a spatula for 15 min or until the Kaiser test was negative. The solution was filtered by vacuum and washed with DMF (3 × 5 mL). The deprotection protocol with TFA was repeated followed by the addition of Boc-L-Phe-OH (530 mg, 2 mmol, 2 equiv) using the elongation protocol described above. The deprotection protocol with TFA was repeated followed by the addition of Boc-L-Val-OH (414 mg, 2 mmol, 2 equiv) with the elongation protocol. After drying, the resin was transferred to a 100 mL oven-dried flask. The reaction vessel was evacuated and backfilled with nitrogen and this process repeated for a total of three times. After the reaction mixture was cooled to 0 °C, TFA (30 mL) was added followed by the addition of thioanisole (3 mL, 25 mmol, 25 equiv), 1,2-ethanedithiol (1.5 mL, 18 mmol, 18 equiv), and trifluoromethanesulfonic acid (3 mL, 34 mmol, 34 equiv). The reaction mixture was warmed to rt over the course of 2 h. At this time, the mixture was filtered and the solid was washed with TFA (2 × 3 mL). To the filtrate was added Et_2_O (100 mL) and extracted with DI H_2_O (3 × 25 mL). The combined aqueous layers were washed with Et_2_O (2 × 50 mL), basified with conc. aqueous NH_4_OH to pH 4 and then concentrated in *vacuo*. The crude residue was partially purified on a CombiFlash^®^ Rf+ equipped with a 50 g RediSep Rf Gold C18Aq column. Acetonitrile and 0.1% trifluoroacetic acid in H_2_O were the mobile phases, and a gradient of 0-100% aq. MeCN was applied over 15 min at 40 mL/min flow rate. The fractions containing **S4** was repurified with a XBridge C18 column (250 × 10 mm, 5 μM particle size). Acetonitrile and 0.1% formic acid in H_2_O were the mobile phases, and a gradient of 5-40% aq. MeCN was applied over 20 min at 5 mL/min flow rate. **S4** was obtained as a white crystalline solid: ^1^H NMR (600 MHz, D_2_O) δ 7.41 (t, J = 7.6 Hz, 2H), 7.33 (m, 3H), 5.20 (s, 1H), 4.73 – 4.70 (m, 1H) 4.37 (q, J = 7.1 Hz, 1H), 3.51 (bs, J = 9.2 Hz, 1H), 3.39 (d, J = 14.8 Hz, 1H), 3.30 (d, J = 14.8 Hz, 1H), 3.21 (dd, J = 13.9, 6.7 Hz, 1H), 3.07 (dd, J = 14.0, 8.8 Hz, 1H), 2.11 – 2.02 (m, 1H), 1.38 (d, J = 7.2 Hz, 3H), 0.95 (d, J = 6.9 Hz, 3H), 0.92 (d, J = 6.8 Hz, 3H); HRMS (ESI) calcd for[C_21_H_31_N_4_O_7_S]^+^ (M+H)^+^: m/z 483.1913 found 483.1898.

**Extended Data Figure 1.**
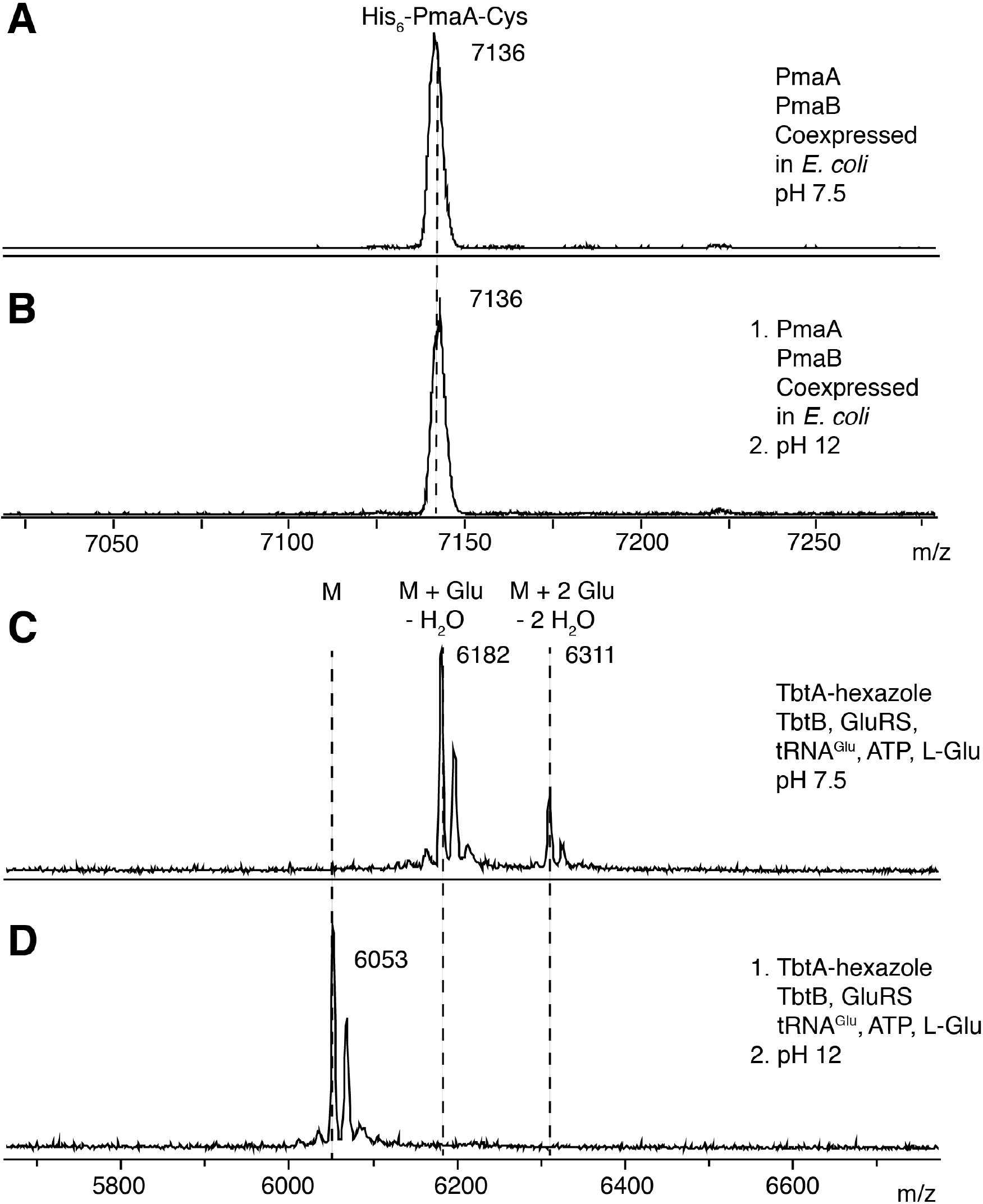
Attempted hydrolysis reaction of PmaA-Cys prepared by co-expression of PmaA and PmaB in *E. coli*. (**a**) pH 7.5 and (**b**) pH 12. The hydrolysis of glutamylated-TbtA hexazole prepared by TbtB and Glu-tRNA^Glu^ is shown as a positive control (**c**) pH 7.5 and (**d**) pH 12. Glutamylated-TbtA hexazole was prepared according to Hudson and co-workers^6^.

**Extended Data Figure 2.**
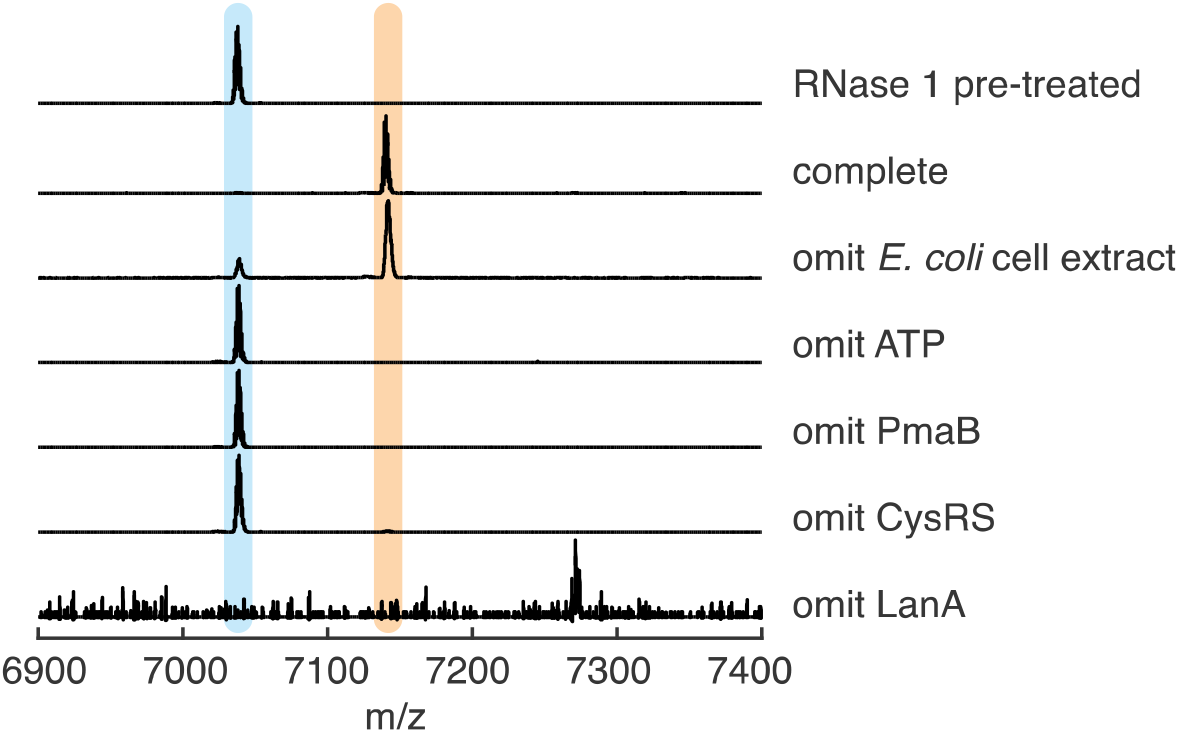
In vitro cysteine addition by PmaB to the precursor peptide PmaA. Peptides were monitored by MALDI-TOF MS. His-tagged proteins and peptides were expressed and purified by IMAC separately. All reactions were incubated at 30 °C for 1 h The complete reaction contains 5 mM ATP, 10 mM Cys, 0.5 μM PmaB, 20 μM PmaA, 0.5 μM CysRS, 5 mM TCEP, 1x assay buffer (50 mM HEPES pH 7.5, 10 mM MgCl_2_, 100 mM NaCl), and a 1:20 dilution of *E. coli* cell extract (see Methods). Then various components were omitted from the assays as indicated or RNase was added prior to initiating the reaction (top mass spectrum).

**Extended Data Figure 3.**
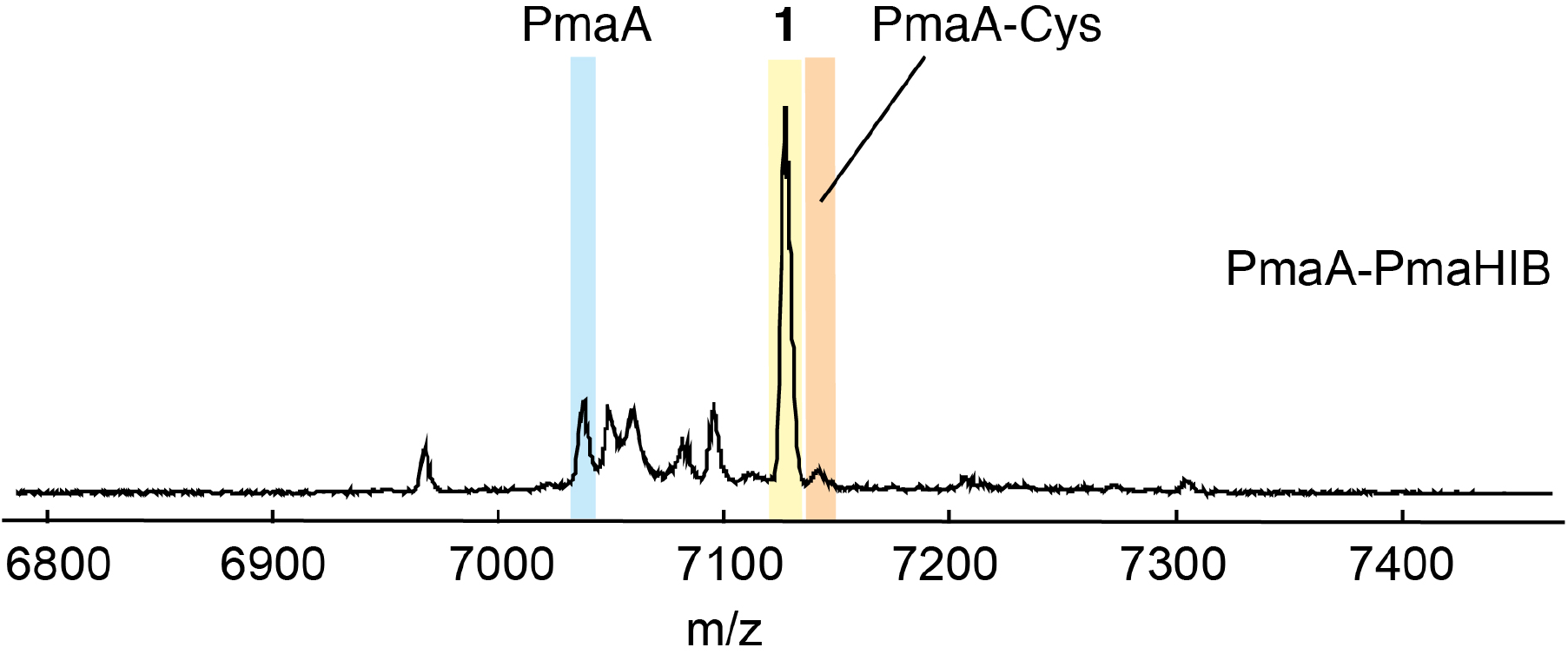
MALDI-TOF mass spectrum of PmaA co-expressed with PmaHIB in *E. coli*.

**Extended Data Figure 4.**
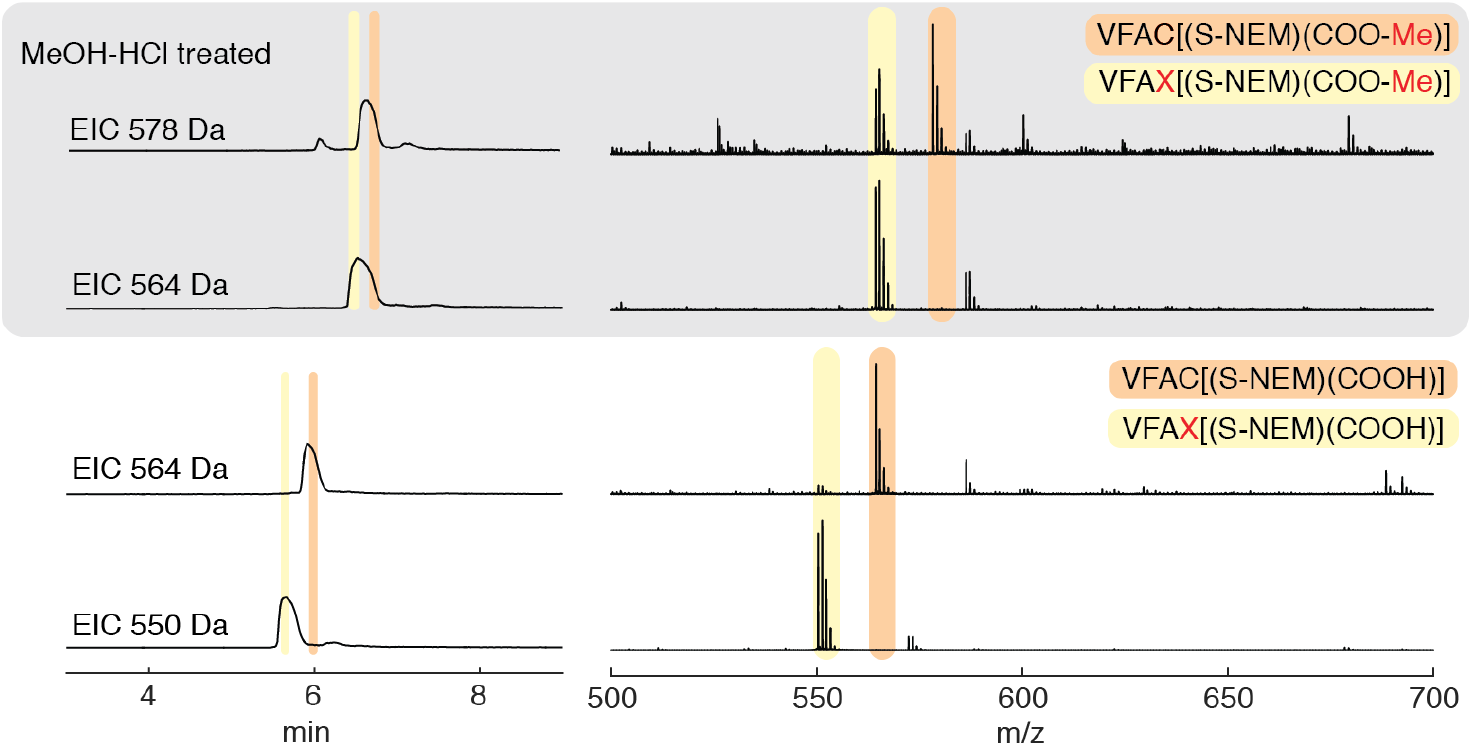
Esterification of PmaA-Cys and the PmaHI-modified peptide following consecutive NEM and trypsin treatment demonstrate preservation of the carboxylate group in the tetrapeptide VFAX. Shaded panel at the top represent experiments that were MeOH/HCl treated, bottom panel are controls that were not MeOH/HCl treated. Extracted ion chromatograms are shown (left) with the corresponding mass spectra of the peaks as marked (right). The observed methylation and NEM-alkylation illustrate that VFAX still contains thiol and carboxylate groups.

**Extended Data Figure 5.**
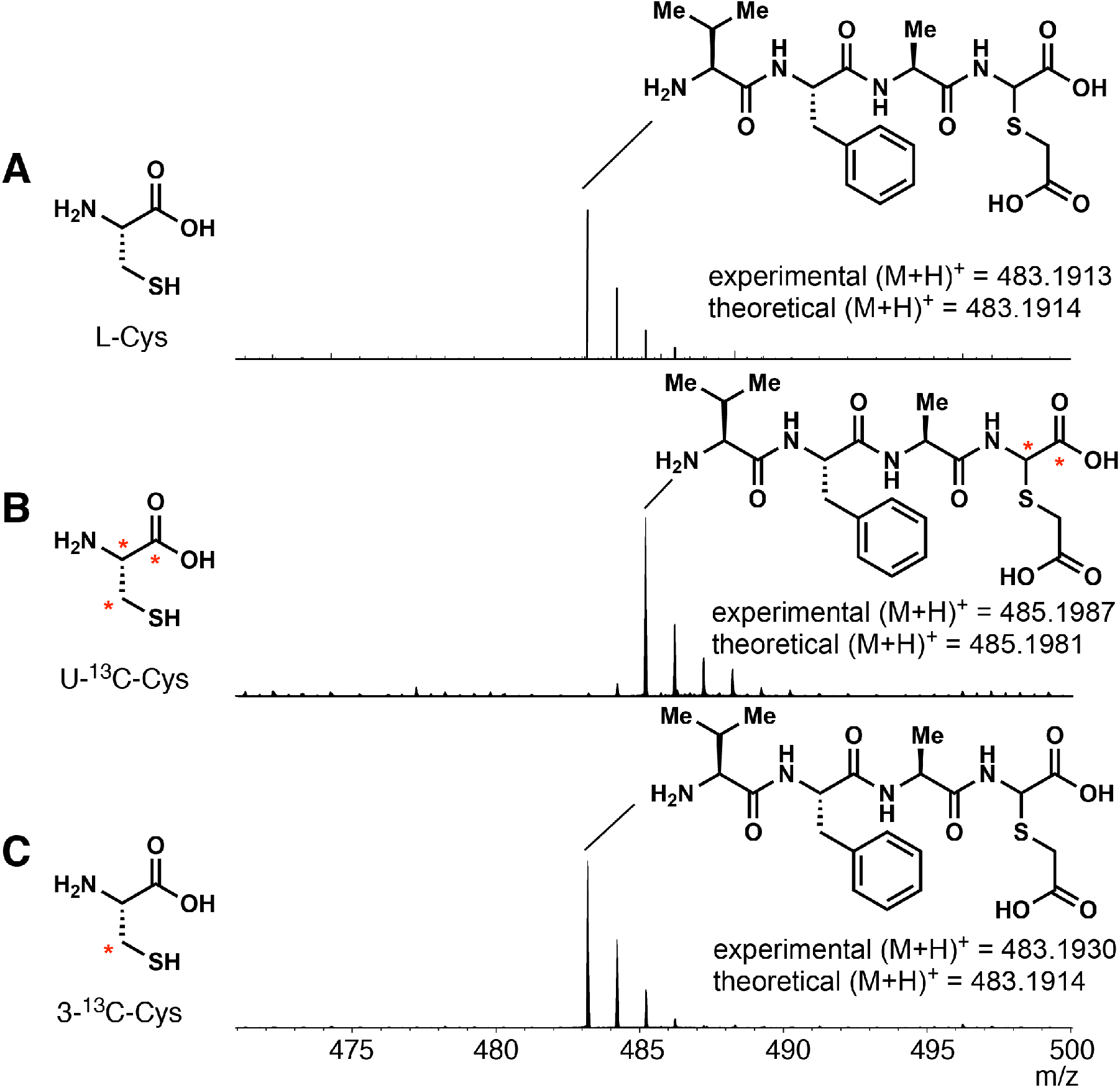
LC-MS spectra (normalized intensity) of trypsin-digested PmaA-Cys modified by PmaHI. Peptides were prepared by co-expression of His_6_-PmaA-Cys and PmaHI in a cysteine auxotrophic strain of *E. coli* grown with (**a**) unlabeled L-cysteine, (**b**) U-^13^C-L-cysteine, and (**c**) 3-^13^C-L-cysteine. Peptides were derivatized with iodoacetic acid prior to analysis.

**Extended Data Figure 6.**
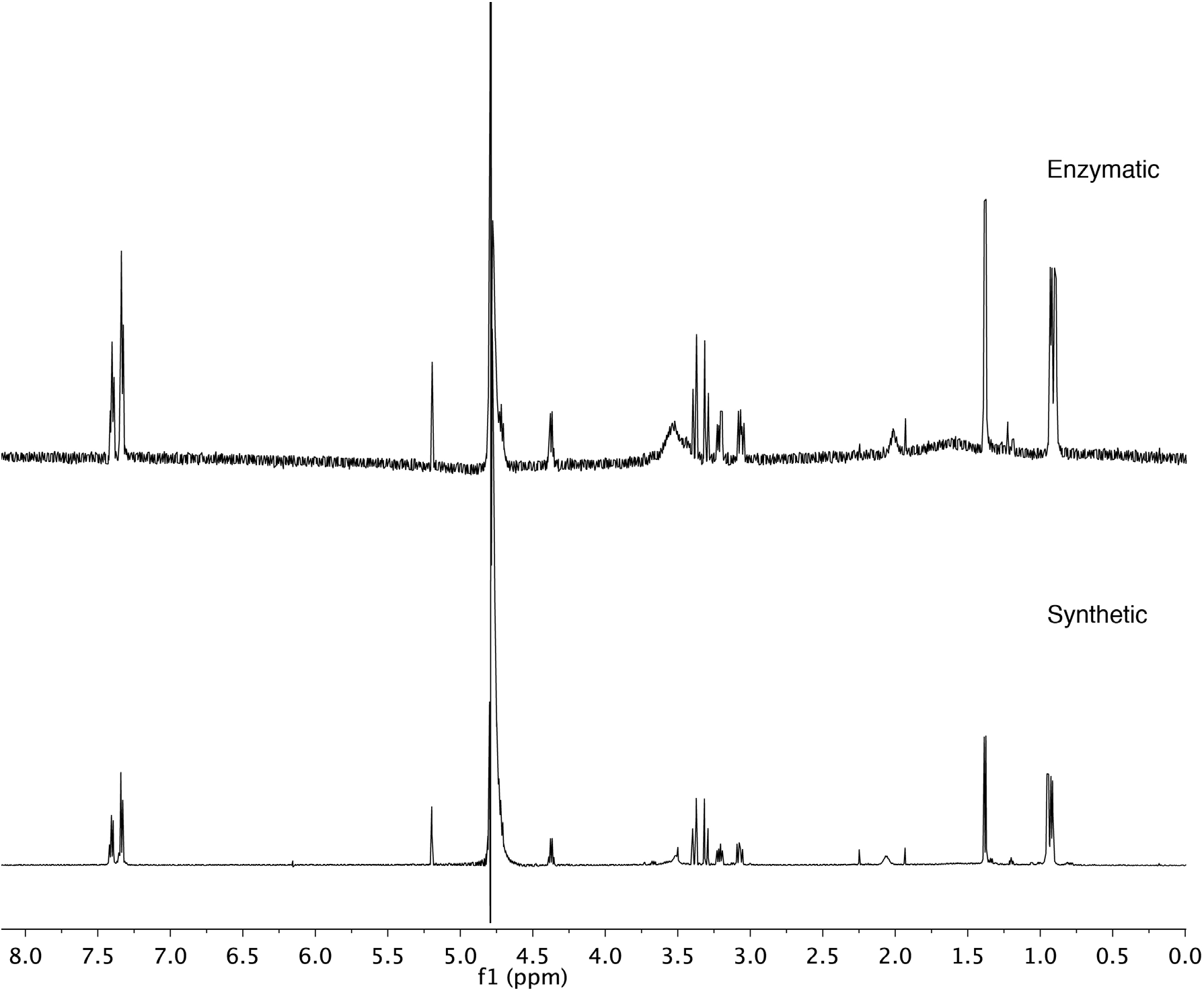
^1^H NMR spectra of VFA-thiaGlu. Top: ^1^H NMR spectrum of VFA-thiaGlu prepared by coexpression of PmaAHIB, subsequent purification and modification by iodoacetic acid, and trypsin digestion. Bottom: ^1^H NMR spectrum of one of the diastereomers of VFA-thiaGlu prepared by chemical synthesis. The other, minor diastereomer could not be obtained in pure form. The signal at 5.2 ppm corresponding to the thioaminal proton is diagnostic.

**Extended Data Figure 7.**
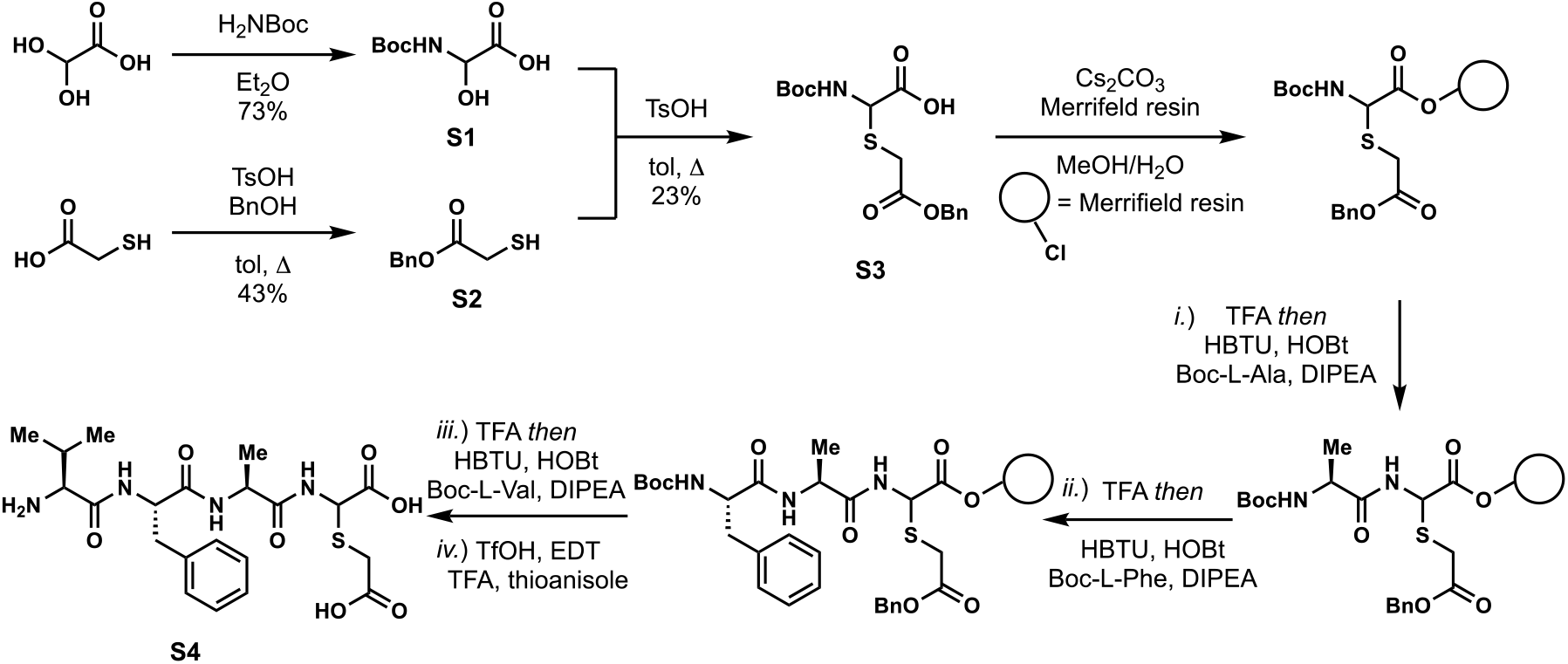
Chemical synthesis of VFA-thiaGlu tetrapeptide.

**Extended Data Figure 8.**
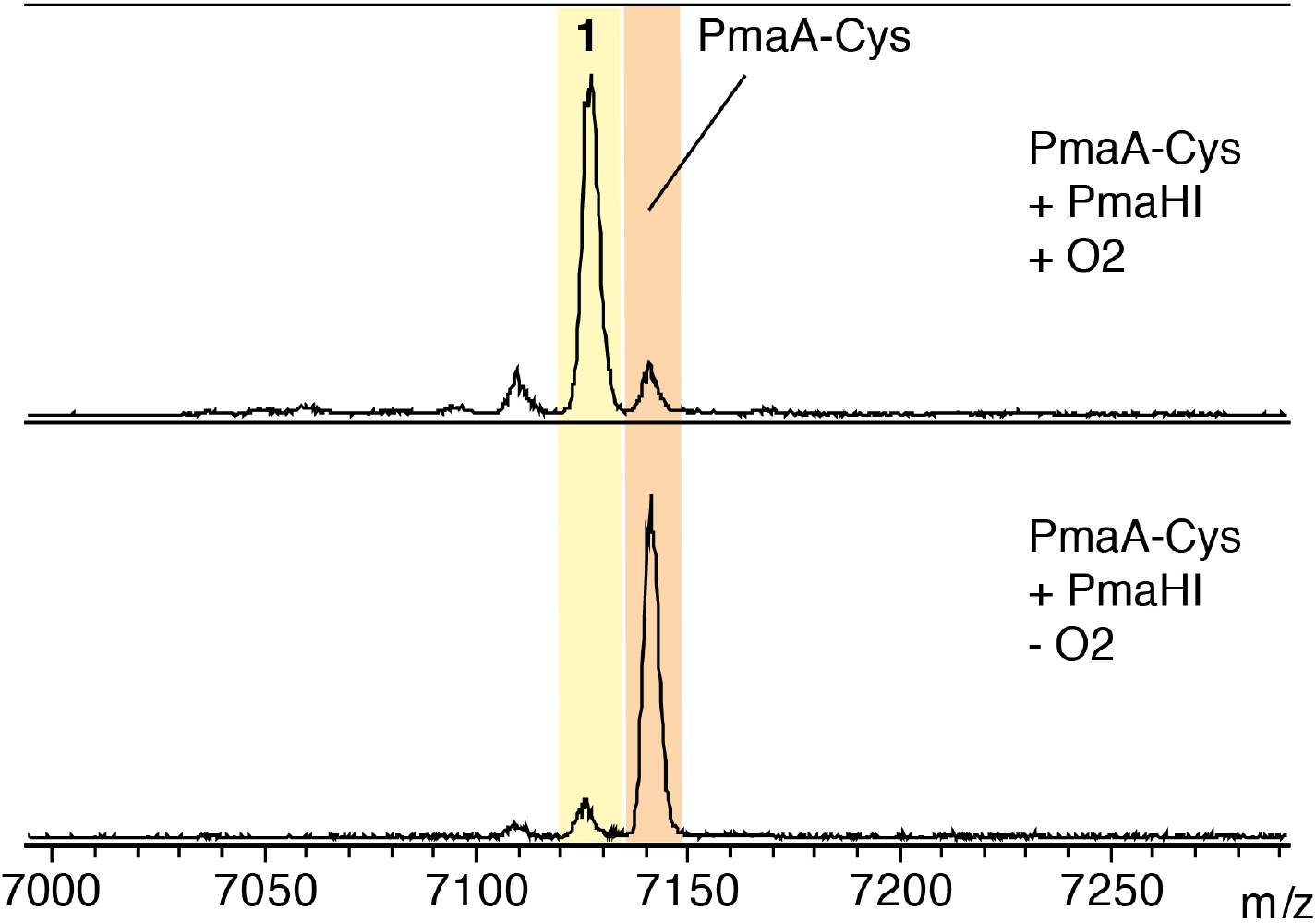
Attempted modification of PmaA-Cys by PmaHI in the absence of oxygen. PmaA-Cys (100 μM) was added PmaHI (10 μM) in phosphate buffer (0.2 mL, 50 mM Na_2_HPO_4_, 300 mM NaCl, 10% glycerol, pH 7.6). The reaction mixture was kept open to air (top spectrum) or in an anaerobic chamber (bottom spectrum) at 24 °C for 1 h upon which it was analyzed by MALDI-TOF MS.

**Extended Data Figure 9.**
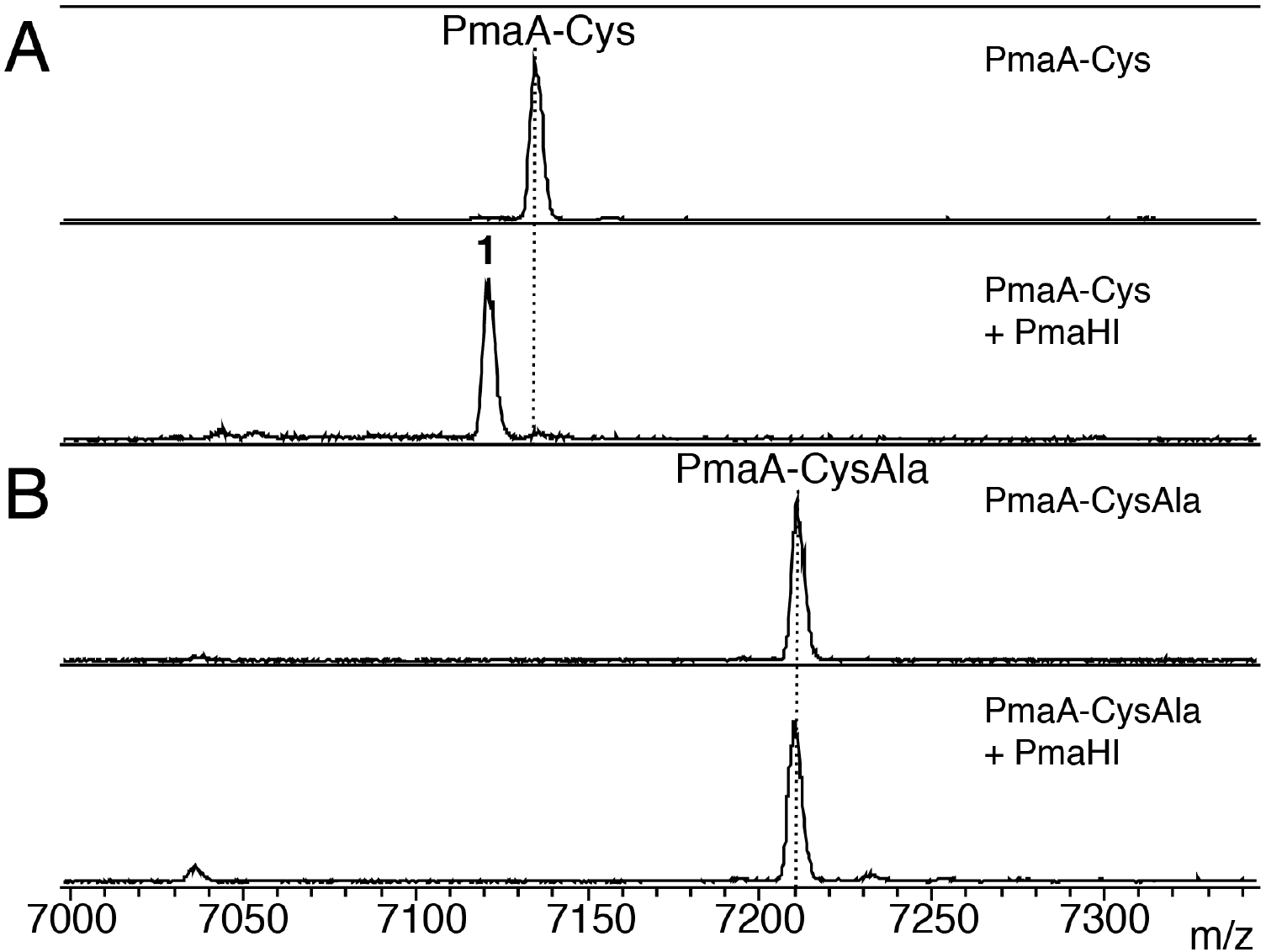
PmaHI requires the carboxylate of the C-terminal cysteine of PmaA-Cys for activity. **a**, MALDI-TOF MS spectrum of PmaA-Cys (100 μM) before and after reaction with PmaHI (20 μM) in phosphate buffer (50 mM Na_2_HPO_4_, 300 mM NaCl, 10% glycerol, pH 7.5). Compound **1** (−14 Da) was observed as the major product along with complete consumption of PmaA-Cys. **b**, MALDI-TOF MS spectrum of PmaA-CysAla (100 μM) before and after reaction with PmaHI (20 μM) in phosphate buffer. No reaction of PmaA-CysAla was detected after 16 h.

**Extended Data Figure 10.**
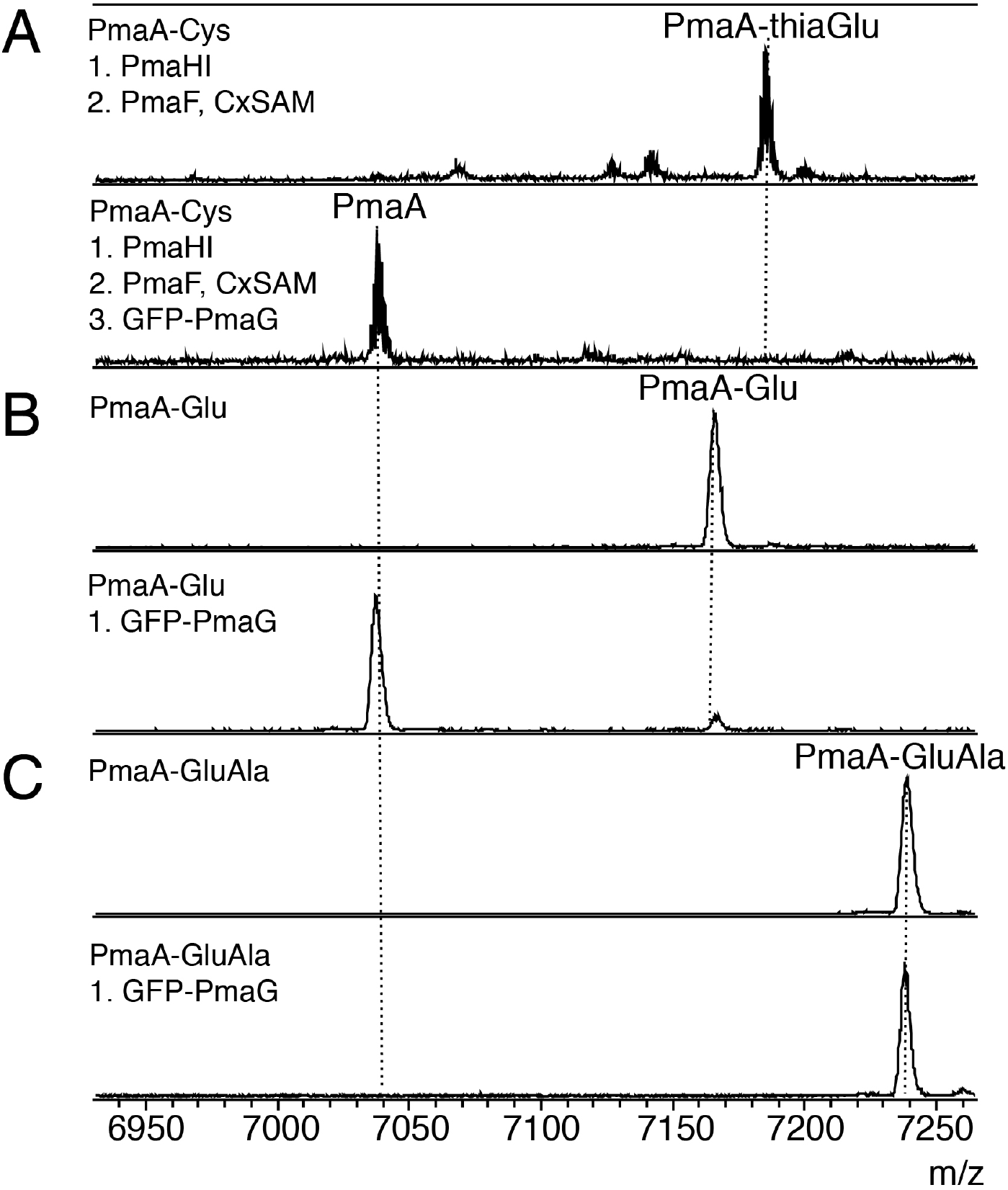
PmaG cleaves off a single amino acid from the C-terminus of the PmaA-3thiaGlu. **a**, MALDI-TOF MS spectrum of PmaA-thiaGlu (**2**) before and after reaction with GFP-PmaG cell lysate in Tris buffer (20 mM Tris, 250 mM NaCl, 10% glycerol, pH 7.5). **b**, MALDI-TOF MS spectrum of PmaA-Glu before and after reaction with GFP-PmaG cell lysate in Tris buffer. **c**, MALDI-TOF MS spectrum of PmaA-GluAla before and after reaction with GFP-PmaG cell lysate in Tris buffer.

**Extended Data Figure 11.**
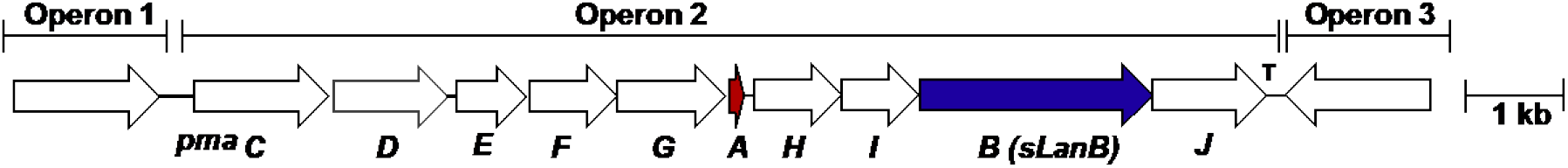
Operon predictions for *pma*. Operon predictions for the cluster in *P. syringae* pv. maculicola ES4326 based on FGENESB^29, 33^.

**Extended Data Figure 12.**
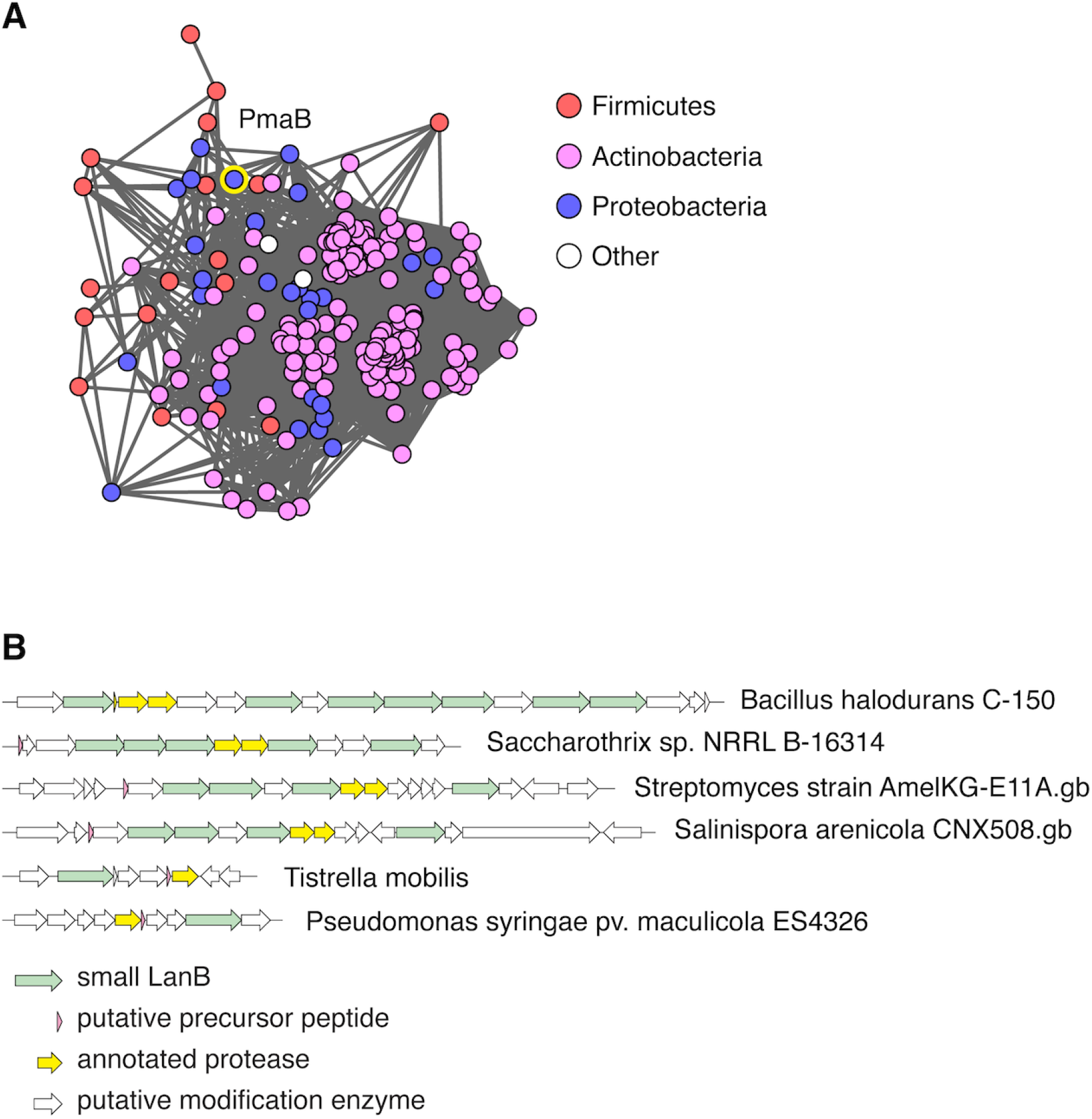
Small LanB enzymes are diverse and are found in many gene contexts. **a**, Sequence similarity network^34^ of small LanB enzymes, annotated by phylogeny. Nodes (circles, N=214) represent sequences clustered at 40% sequence identity. Lines (N=12279) represent connections drawn at an alignment score E value of - 20. Network was generated from sequences in PFAM group PF04738 at an E value of −5 and trimmed to remove clusters containing full length LanB enzymes at an E value of - 20. Colors are assigned based on phylum level annotation. The node containing PmaB is highlighted in yellow and has 10 neighbors at the cut-off. **b**, Example biosynthetic gene clusters containing *pmaB*-related genes. Five examples were chosen for each analysis based on diversity of the cluster architecture; the *pma* cluster is shown at the bottom for comparison.

**Extended Data Figure 13.**
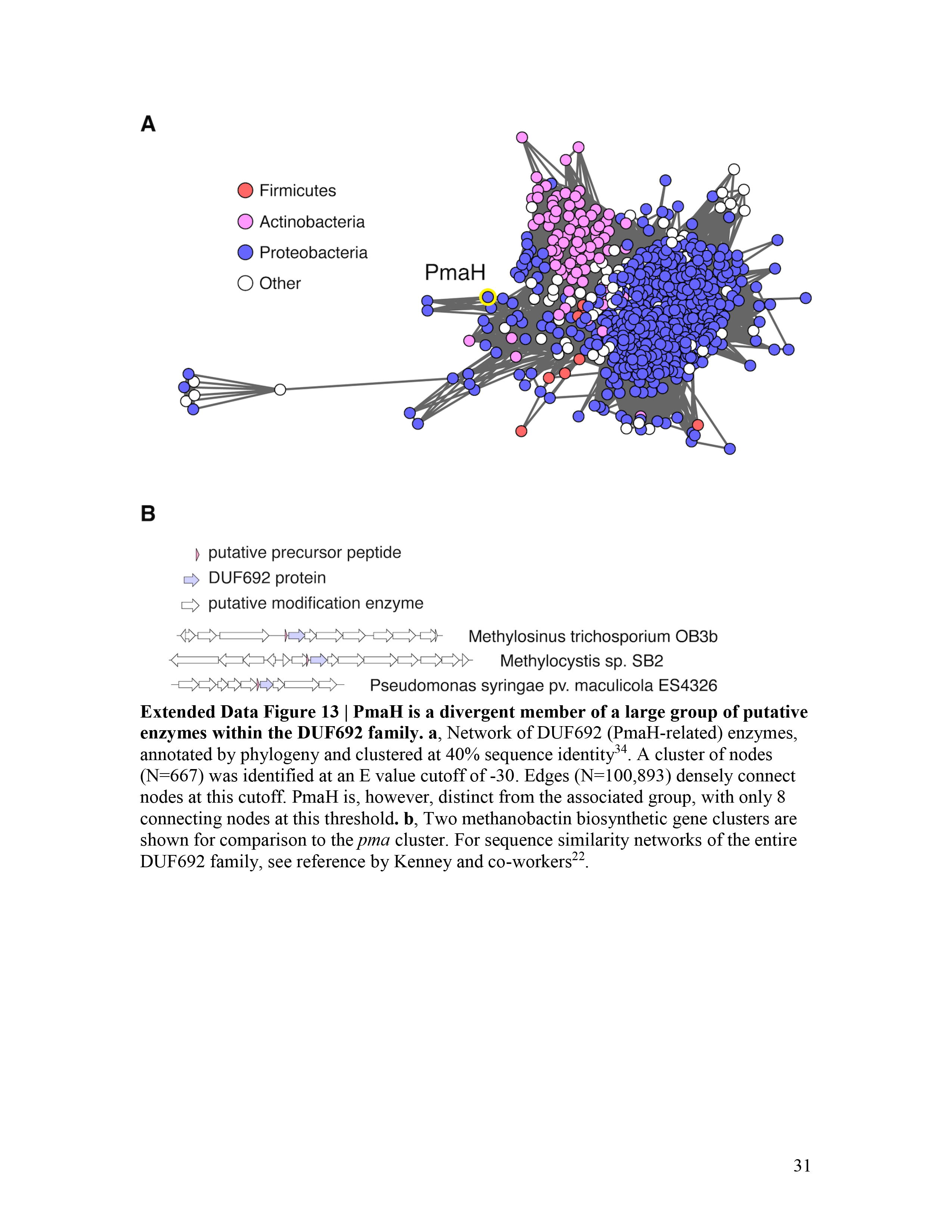
PmaH is a divergent member of a large group of putative enzymes within the DUF692 family. **a**, Network of DUF692 (PmaH-related) enzymes, annotated by phylogeny and clustered at 40% sequence identity^34^. A cluster of nodes (N=667) was identified at an E value cutoff of −30. Edges (N=100,893) densely connect nodes at this cutoff. PmaH is, however, distinct from the associated group, with only 8 connecting nodes at this threshold. **b**, Two methanobactin biosynthetic gene clusters are shown for comparison to the *pma* cluster. For sequence similarity networks of the entire DUF692 family, see reference by Kenney and co-workers^22^.

**Extended Data Figure 14.**
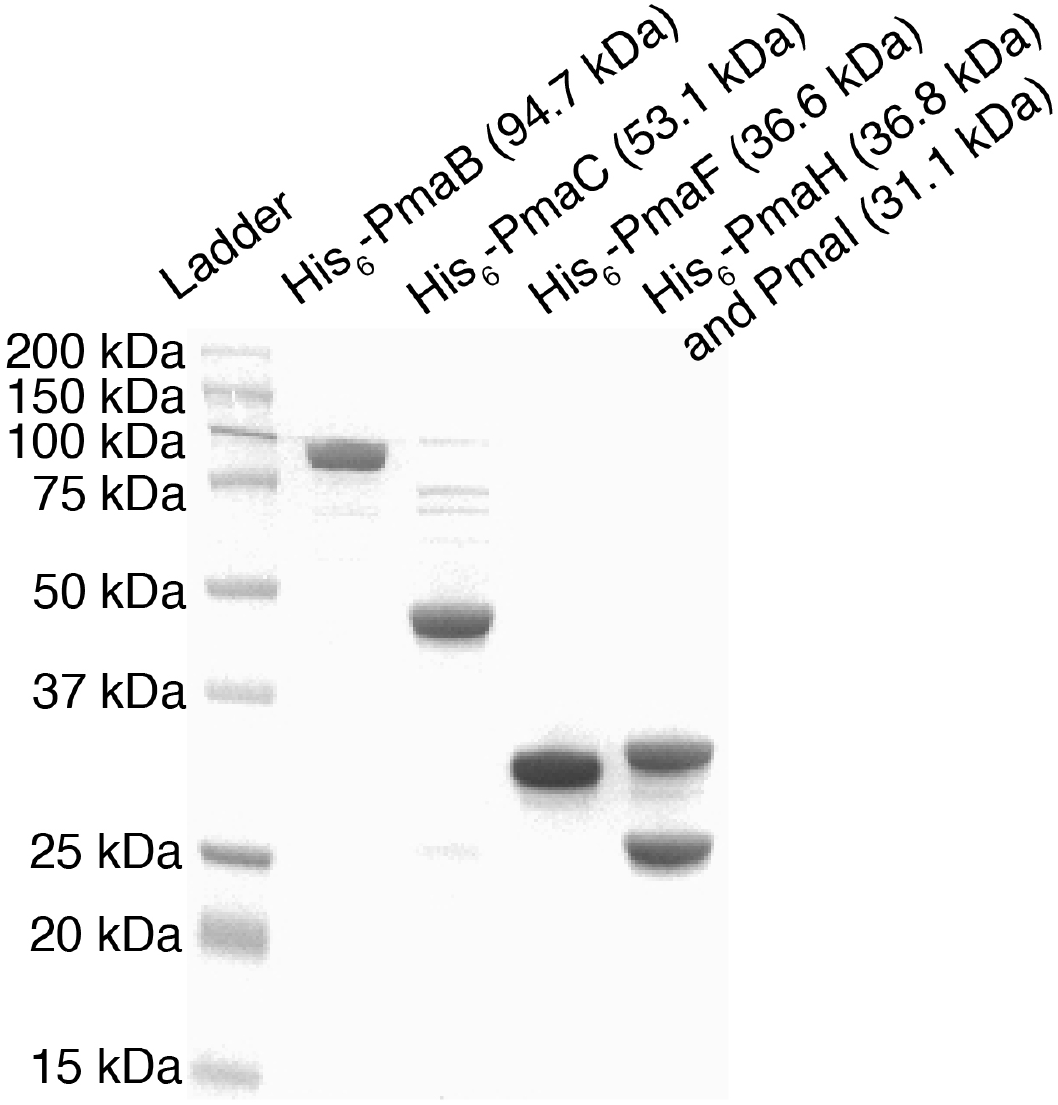
SDS-PAGE of all proteins used in this study.

**Extended Data Table 1.**
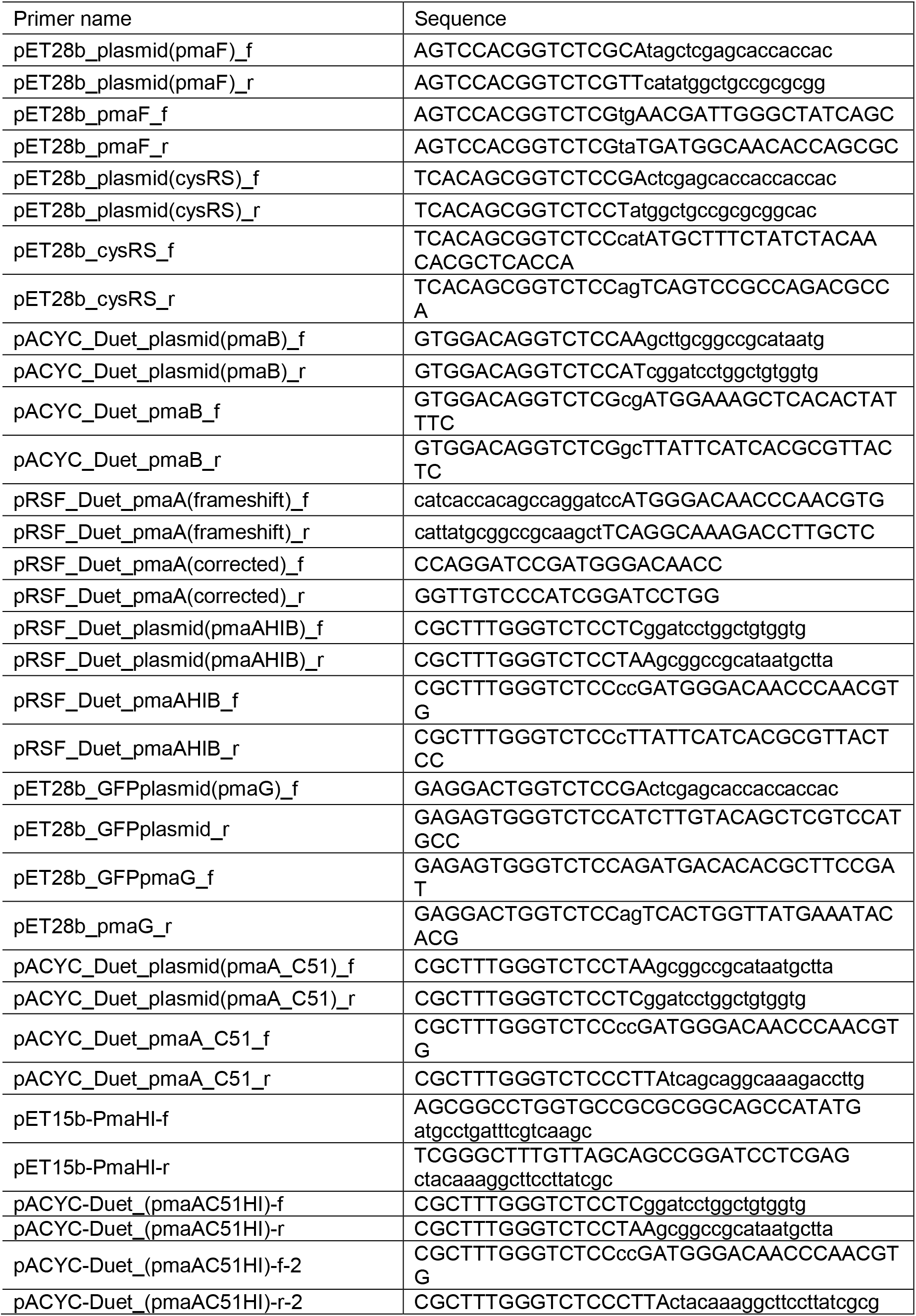
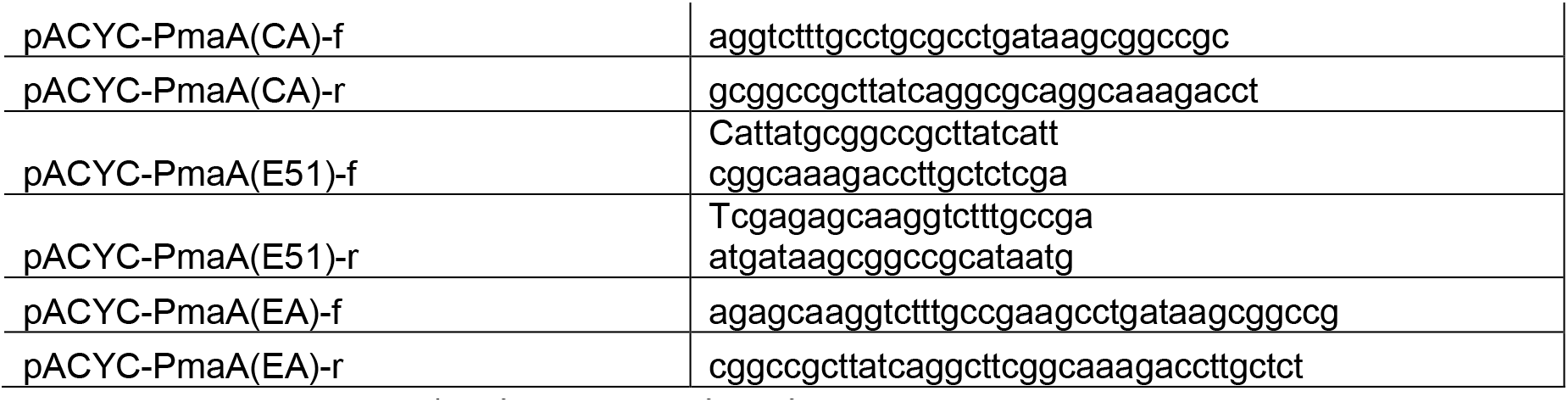
Primers used in this study.

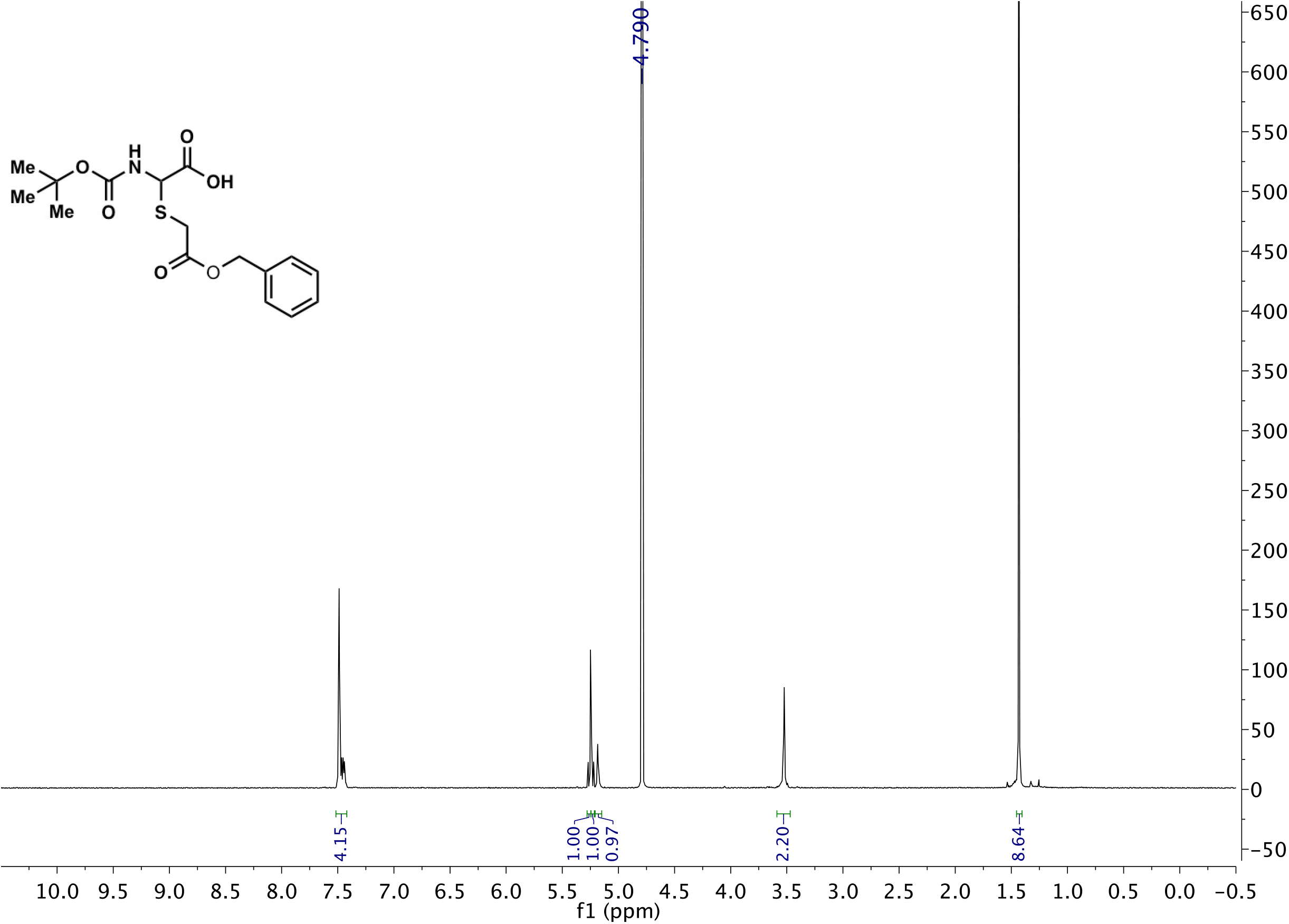

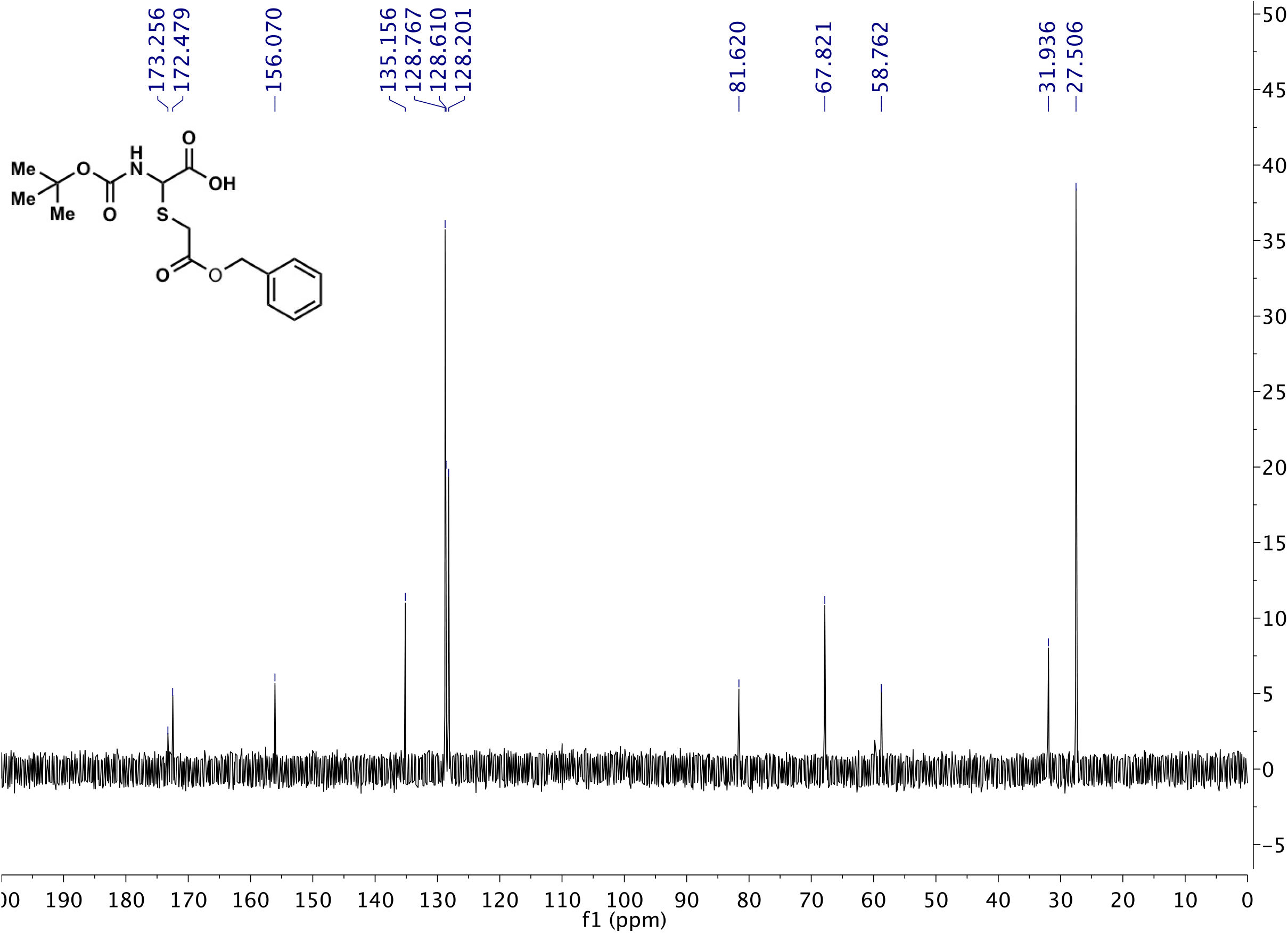

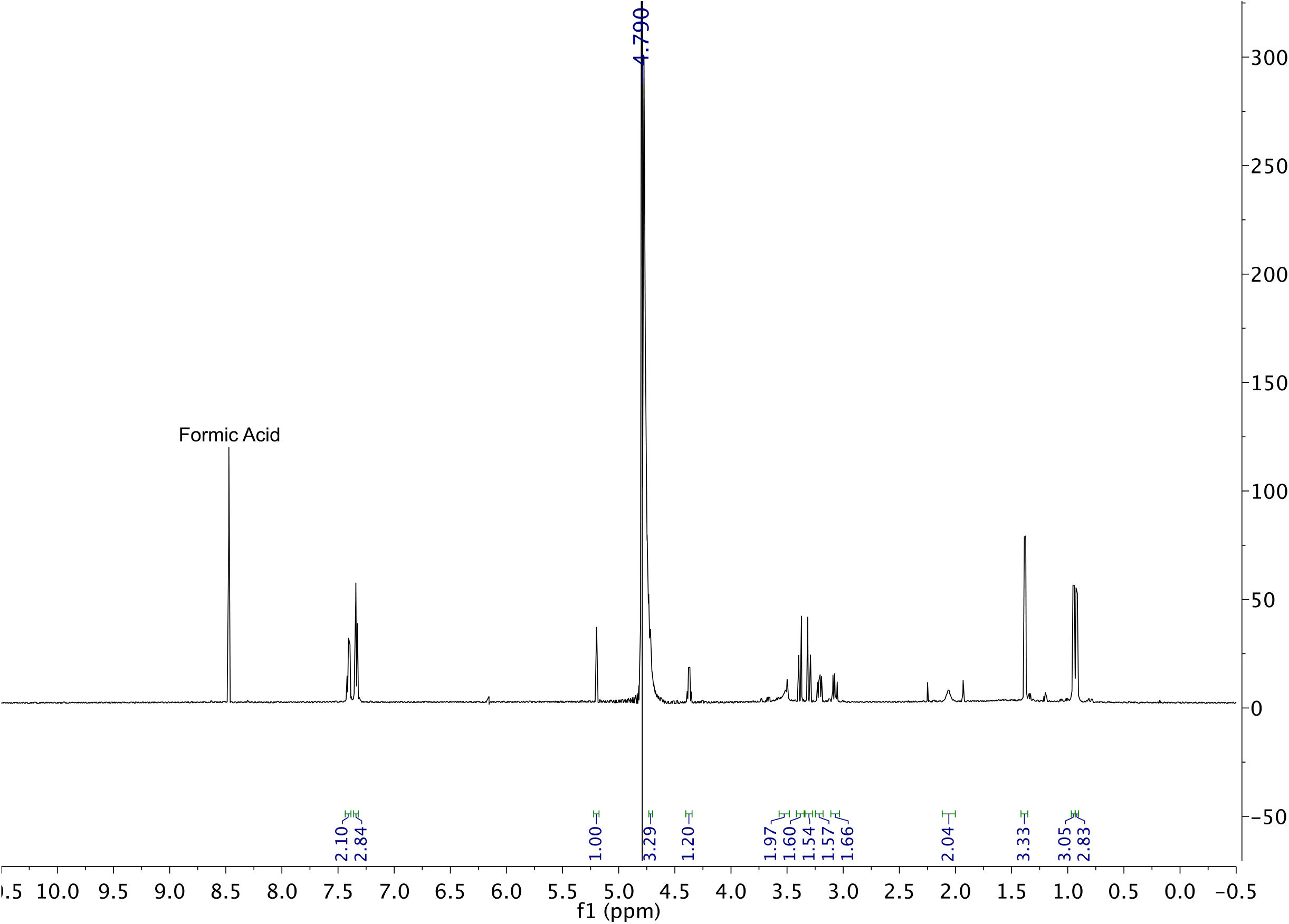

